# Structural basis for the allosteric regulation and dynamic assembly of DNMT3B

**DOI:** 10.1101/2023.09.09.556992

**Authors:** Jiuwei Lu, Jian Fang, Hongtao Zhu, Kimberly Lu Liang, Nelli Khudaverdyan, Jikui Song

**Affiliations:** Department of Biochemistry, University of California, Riverside, CA 92521, USA; Vollum Institute, Oregon Health and Science University, Portland, OR, USA; Diamond Bar High School, Diamon Bar, CA 91765, USA

## Abstract

Oligomerization of DNMT3B, a mammalian *de novo* DNA methyltransferase, critically regulates its chromatin targeting and DNA methylation activities. However, how the N-terminal PWWP and ADD domains interplay with the C-terminal methyltransferase (MTase) domain in regulating the dynamic assembly of DNMT3B remains unclear. Here, we report the cryo-EM structure of DNMT3B under various oligomerization states. The ADD domain of DNMT3B interacts with the MTase domain to form an autoinhibitory conformation, resembling the previously observed DNMT3A autoinhibition. Our combined structural and biochemical study further identifies a role for the PWWP domain and its associated ICF mutation in the allosteric regulation of DNMT3B tetramer, and a differential functional impact on DNMT3B by potential ADD-H3K4me0 and PWWP-H3K36me3 bindings. In addition, our comparative structural analysis reveals a coupling between DNMT3B oligomerization and folding of its substrate-binding sites. Together, this study provides mechanistic insights into the allosteric regulation and dynamic assembly of DNMT3B.

## Introduction

DNA methylation is an evolutionarily conserved gene regulatory mechanism that critically influences chromatin structure and function (1). In mammals, DNA methylation predominantly occurs at the C-5 position of cytosines within CpG dinucleotides, accounting for ∼70% of total CpG sites in the genome (2). Mammalian DNA methylation is mainly established by *de novo* DNA methyltransferases DNMT3A and DNMT3B during gametogenesis and embryogenesis (3). A single knockout (KO) of DNMT3A or DNMT3B resulted in embryonic or postnatal mortality, testifying that both enzymes are essential for development (3). Along the line, mutations of DNMT3A have been associated with hematological disorders (e.g. Acute Myeloid Leukemia) (4) and growth anomalies (e.g. Tatton-Brown-Rahman syndrome) (5), whereas DNMT3B mutants account for over 60% of patients with immunodeficiency, centromeric region instability, and facial anomalies (ICF) syndrome (3,6,7). Development of therapeutic strategies for these diseases necessitate a detailed understanding of the functional regulation of DNMT3A and DNMT3B.

DNMT3A and DNMT3B are closely related in sequence, each containing a C-terminal methyltransferase (MTase) domain preceded by a Pro-Trp-Trp-Pro (PWWP) domain and ATRX-DNMT3-DNMT3L (ADD) domain (8). It has been established that the DNA methylation activities and chromatin targeting of DNMT3A and DNMT3B are controlled by mechanisms of homo-oligomerization (9–12) and/or hetero-oligomerization (13–18). Previous studies have indicated the tetrameric assembly as the most stable and active form of DNMT3A and DNMT3B *in vitro* (9,10,12,19), which may transit into a filamentous, high-order oligomeric assembly (denoted as macro-oligomer thereafter) in the presence of DNA (12). Disruption of DNMT3A or DNMT3B oligomerization by the interface mutations was found to impair their DNA methylation activities and/or heterochromatin targeting in cells (9–11,16,17,20). Conversely, DNMT3A R882H, a hot-spot mutation in AML (21), was shown by several studies to promote macro-oligomer formation of DNMT3A (22,23), which compromises the DNA methylation activity of DNMT3A in stemness-related promoters, thereby contributing to leukemogenesis (23,24).

DNMT3A- and DNMT3B-mediated *de novo* DNA methylation is both subject to regulation by DNMT3L (13–15), albeit to a different extent (3,13,15,25), with the latter directly interacting with DNMT3A or DNMT3B to stimulate its DNA methylation activity (for DNMT3A and DNMT3B) (14,26–29) or stability (for DNMT3A) in cells (18). A pioneering structural study of the C-terminal domains of DNMT3A-DNMT3L reveals a tetrameric assembly, mediated by a polar interface (a.k.a. RD interface) for the DNMT3A homodimerization, and a nonpolar interface (a.k.a. FF interface) for the association of each DNMT3A monomer with DNMT3L (16). Subsequent studies from others and us demonstrated that the complexes of the C-terminal domains of DNMT3B-DNMT3L and DNMT3A-DNMT3L adopt a similar conformation for substrate interactions, with the RD interface directly involved in DNA binding (30–33). In addition, an intramolecular interaction between the RD interface and a loop from the target recognition domain (TRD) regulates the disorder-to-order transition of the latter upon DNA binding, which is critical for the CpG-specific recognition by DNMT3B/DNMT3A (30,31,33).

The functions of DNMT3A and DNMT3B are further regulated by their N-terminal domains (8,34–36). The DNMT3A ADD domain, a reader module of histone H3K4me0 mark (37,38), interacts with the MTase domain to form an autoinhibitory state, which can be relieved by the H3K4me0 binding, thereby coupling H3K4me0 binding with enzymatic activation (39). The PWWP domains of DNMT3A and DNMT3B were shown to bind to both DNA and H3 lysine 36 di- or tri-methylation (H3K36me2/3) (40–42), which is essential for targeting DNMT3A and DNMT3B to heterochromatin (40,41,43) and/or gene body (44,45). To date, no structure of DNMT3A and DNMT3B in the context of all three functional domains has been reported. How the N-terminal PWWP and ADD domains are coordinated in regulating the activity and assembly of DNMT3A and DNMT3B has yet to be characterized.

To investigate the functional regulation of DNMT3B, we determined the single-particle cryogenic electron microscopy (cryo-EM) structure of a DNMT3B fragment comprised of the PWWP, ADD and MTase domains. Analysis of the DNMT3B assemblies reveals the molecular basis for DNMT3B homotetramers and ascending/descending oligomeric states. The association of DNMT3B ADD domain with the MTase domain results in an autoinhibitory conformation that is similar but distinct from the previously observed autoinhibitory state of DNMT3A-DNMT3L complex. Structural analysis, combined with biochemical and *in vitro* DNA methylation assays, further reveals an interaction of the PWWP domain with ADD and MTase domains, which reinforces the allosteric regulation of DNMT3B homotetramer, and underpins a differential impact of histone H3K4me0 and H3K36me3 bindings on DNMT3B activity. In addition, structural analysis of individual DNMT3B subunits reveals a coupling between the integrity of the RD interface and the folding of the TRD and the cofactor-binding site, suggesting an oligomerization-coupled substrate binding activity. Together, these observations provide a structural basis for both the allosteric regulation and dynamic assembly of DNMT3B, with important implications for the functional regulation of DNMT3B in development and diseases.

## Materials and Methods

### Protein expression and purification

DNA encoding a fragment of human DNMT3B, PWWP-ADD-MTase (residues 206-853), ADD-MTase (residues 396-853) or MTase (residues 569-853), was inserted into an in-house bacterial expression vector, preceded by an N-terminal hexahistidine (His6)-MBP tag and a TEV cleavage site. BL21(DE3) RIL cells harboring the resulting plasmids were induced by addition of 0.067mM isopropyl β-D-1-thiogalactopyranoside (IPTG) with 50 µM ZnCl2 when the cell density reached A600 of 1.0 and continued to grow at 16 °C overnight. The cells were harvested and lysed in buffer containing 50 mM Tris-HCl (pH 8.0), 1 M NaCl, 25 mM Imidazole, 10% glycerol, 10 µg/mL DNase I and 1 mM PMSF. Subsequently, the fusion proteins were purified through a nickel column, followed by removal of (His6)-MBP tag by TEV cleavage, ion exchange chromatography using a HiTrap Heparin HP column (GE Healthcare) and further purification through size-exclusion chromatography on a Hiload 16/600 Superdex 200 column (GE Healthcare) in buffer containing 20 mM Hepes-NaOH (pH 7.2), 250 mM NaCl, 5% glycerol and 5 mM DTT. The purified protein samples were concentrated and stored at -80 °C before use. The mutants were constructed by site-directed mutagenesis and purified in the same way as described above.

DNMT3B PWWP (residues 206-355), ADD (residues 417-555) and mutants were cloned into a modified pRSF vector preceded by an N-terminal His6-Sumo tag and a ULP1 cleavage site. The BL21(DE3) RIL cells were induced with 0.4 mM IPTG with 50 µM ZnCl2 when A600 reached 0.8 and continued to grow at 16 °C overnight. Similar to the purification of the long DNMT3B fragment (residues 206-853) described above, the proteins were purified sequentially via nickel affinity chromatography, removal of His6-Sumo tag by ULP1 cleavage, ion exchange chromatography using a HiTrap Heparin HP column (GE Healthcare) for the PWWP domain or a HiTrap Q HP column (GE Healthcare) for the ADD domain, and size-exclusion chromatography using a Hiload 16/600 Superdex 75 column (GE Healthcare). The purified protein sample was stored at -80 °C before use.

### Size-exclusion chromatography analysis of DNMT3B (206-853)

Size-exclusion chromatography analysis of purified DNMT3B (206-853) protein, either untreated or subject to chemical crosslinking by 0.02% Glutaraldehyde for 15 minutes, was carried out on a Superose 6 increase 10/300 column (GE Healthcare) pre-equilibrated with buffer containing 20 mM Tris-HCl (pH 7.2), 250 mM NaCl, 5% Glycerol and 5 mM DTT. The elution volumes for the molecular weight markers and full-length Flag-tagged DNMT3B on the same Superose 6 increase column, as determined in our recent study (9), were used for comparison.

### ITC measurements

Human DNMT3B PWWP and ADD were dialyzed at 4°C overnight against a buffer containing 20 mM Tris-HCl (pH 7.5), 100 mM NaCl and 1 mM β-mercaptoethanol. The final concentrations, determined based on ultraviolet absorption at 280 nm, were 0.2 mM for ADD and 2 mM for PWWP and mutations. A MicroCal iTC200 system (GE Healthcare) was used to conduct the ITC measurements. A total of 15∼17 injections with a spacing of 180 second and a reference power of 5 μcal/s were performed after the temperature was equilibrated to 20°C. The ITC curves were processed with software Origin (v7.0, MicroCal) using one-site fitting model.

### Cryo-EM sample preparation and data acquisition

The protein sample of human DNMT3B (206-853) was crosslinked on ice for 15 minutes at 1.3 mg/mL with 0.02% Glutaraldehyde, followed by size-exclusion chromatography to remove over-crosslinked aggregation on a Superose 6 increase 10/300 column (GE Healthcare) pre-equilibrated with buffer containing 20 mM Tris-HCl (pH 7.2), 250 mM NaCl, 5% Glycerol and 5 mM DTT. The peak fractions were collected and concentrated to negative-stain electron microscopy for sample optimization.

For cryo-EM sample preparation, an aliquot of 2.5 μL of the above optimized DNMT3B (206-853) at a concentration of approximately 0.5 mg/ml (OD280: 0.8) was applied to a Quantifoil holey carbon grid (Copper,1.2/1.3, 300 mesh), which was glow discharged for 1 min with H2/O2 through Gatan Plasma clean System (SOLARUS). The grid was blotted and plunge-frozen in liquid ethane cooled by liquid nitrogen with a Vitrobot IV (Thermo Fisher) at 4 °C under 100% humidity. The frozen grids were stored in liquid nitrogen before use.

High-resolution cryo-EM data were collected on a Titan Krios electron microscope operating at 300 kV, equipped with a post-GIF Gatan K3 Summit (BioQuantum) direct electron detector (counting mode) in the Stanford-SLAC Cryo-EM center using EPU (v2.7, Thermo fisher). Movies were recorded at a nominal magnification of ×81,000 with a pixel of 1.1 Å with the defocus range of −1.5 and −2.1 μm. Each micrograph was recorded 40 frames at a dose rate of 12.5 *e-*/ Å 2/s and total expose time of 4 s, resulting in a total dose rate of 50 *e−*/Å2.

### Cryo-EM data processing

The cryo-EM data for DNMT3B were processed using cryoSPARC (v3.3.1) (46). The movies were motion-corrected and dose-weighted using the patch-based motion correction module. The contrast transfer function (CTF) of the resulting images, with a pixel size of 1.1 Å, were then subject to patch-based estimation. Automatic particle picking was performed using the TOPAZ method (47). After training and particle picking with TOPAZ, 5,145,319 particles were exacted from 8,970 images, with a down-scaled pixel size of 5.2 Å. After two rounds of clean-up by 2D classifications, those classes with identifiable secondary or tertiary structures, comprised of 1,558,376 particles, were selected. Initial models were then generated by 3D *ab initio* reconstruction followed by one round of heterogenous refinement applying C1 symmetry. The particles associated with the four classes with clearly defined DNMT3B oligomer densities were each re-extracted with a pixel size of 1.1 Å. Another round of 3D classification was performed for each class. Four different classes were generated: 185,721 particles for the tetramer I, 163,475 particles for the tetramer II, 130,219 particles for the trimer, and 169,827 particles for the hexamer. After a final non-uniform refinement in cryoSPARC, the resolution for the tetramer I, tetramer II, trimer and hexamer were at 3.04, 3.12, 3.34 and 3.19 Å, respectively, as given by the Fourier shell correlation criterion (FSC 0.143). CryoSPARC was used to determine local resolution and sharpen the maps.

### Model building

For model building of the DNMT3B tetramer I, the reported crystal structures of the DNMT3A ADD domain (PDB 4U7P) (39), a tetrameric unit of the DNMT3B macro-oligomer (PDB 7V0E) (9) and the DNMT3B PWWP domain (PDB 5CIU) (42) were used to fit into the cryo-EM map of the tetramer I in Chimera X (v1.4) (48). The initial structural model was then subject to iterative model building using Coot (v0.9.6) (49) and real-space refinement using Phenix (v1.8.2) (50). The structural models of the DNMT3B tetramer II, trimer and hexamer were generated by fitting the structural model of the tetramer I into the cryo-EM map of the corresponding assemblies, followed by domain/subunit removal or addition in coot to conform with the cryo-EM density. The final structural models of DNMT3B tetramer II, trimer and hexamer were obtained after iterative model building in coot and real-space refinement in Phenix, as with that for the tetramer I. For all DNMT3B assemblies, a model-map Fourier shell correlation was calculated using the criterion of 0.5.

### DNA methylation assay

A 652-bp DNA containing multiple CpG sites derived from a fragment of pGEX-6P-1 vector (51) was used as substrate. For histone H3-mediated stimulation, The H3(1–15), H3(16–41)K36me3, H3(1–41) and H3(16–41)K36me3 peptides were synthesized from LifeTein LLG, with H3(1–15) containing an additional C-terminal tyrosine for spectroscopic quantification. For DNA methylation reactions, 1 μM DNMT3B PWWP-ADD-MTase (residues 206-853) protein, WT or mutant, was pre-incubated with or without 10 μM histone H3 peptide. A 20-μL reaction mixture contained 1 µg DNA, 0.5 μM DNMT3B protein, 5 µM histone H3 peptide, 0.55 μM S-adenosyl-L-[methyl-3H] methionine (specific activity 58.9 Ci/mmol, PerkinElmer) in 50 mM Tris–HCl (pH 7.2), 250 mM NaCl, 0.05% β-mercaptoethanol, 10% glycerol, 0.01% NP-40 and 200 μg/mL BSA. The DNA methylation assays were carried out in triplicate at 37°C for 30 min. Subsequently, the reaction was quenched by addition of 5 μL of 10 mM AdoMet. For detection, 8 μL of the reaction mixture was spot on Amersham Hybond-XL paper (GE healthcare) and dried out. The paper was then washed with 0.2 M cold ammonium bicarbonate (pH 8.2) (twice), Milli Q water and Ethanol. Subsequently, the paper was air dried and transferred to scintillation vials filled with 3 mL of ScintiVerse cocktail (Thermo fisher). The radioactivity of tritium was measured with a Beckman LS6500 counter.

### Dynamic light scattering (DLS)

Approximately 0.7 mg/mL of WT or mutant DNMT3B PWWP-ADD-MTase, ADD-MTase or MTase fragment was dissolved in buffer containing 20 mM HEPES (pH 7.2), 100 or 250 mM NaCl, and 5 mM DTT. DLS of the samples was measured in a 384-well microwell plate (3820, Corning) using DynaPro Plate Reader II (Wyatt Technology Corporation) at 25 °C. The hydrodynamic radii of the DNMT3B fragments were derived from triplicate measurements using the DYNAMICS software (Wyatt Technology Corporation).

### Thermal shift assay

Each sample mixture contains 1 µM WT or mutant DNMT3B PWWP-ADD-MTase dissolved in 20-µL buffer containing 20 mM HEPES (pH 7.2), 5% glycerol, 250 mM NaCl, 5 mM DTT and 1X GloMelt Dye. The experiment was conducted using a BioRad CFX96 Connect Real-Time PCR detection system, as previously described (52), with the sample plate subjected to stepwise heating from 4°C to 95°C at 0.5°C per increment. Fluorescence intensity was recorded with the excitation and emission wavelength set to 470 nm and 510 nm, respectively. The experiments were performed in triplicate.

## Results

### Assembly overview of DNMT3B

To illustrate how the regulatory domains and the MTase domain of DNMT3B cooperate in controlling its DNA methylation activity and assembly, we characterized the structure of DNMT3B by single-particle cryo-EM approach. We focused on a DNMT3B fragment (residues 206-853: DNMT3B206-853; Fig. 1A) that harbors the PWWP, ADD and MTase domains but can be more manageably purified than full-length DNMT3B. Size-exclusion chromatography analysis of this fragment indicates a similar assembly state as that of full-length DNMT3B purified from mouse embryonic stem cells (Fig. S1), which was dominated by a tetrameric form as characterized previously (9). Analysis of the cryo-EM data led to identification of four assembly states, including two alternative forms of tetramers (tetramer I and tetramer II) that together account for over half of the final particles, a trimeric form, and a hexameric form (Fig. 1B-E and Fig. S2 and Table S1). The structures of tetramer I, tetramer II, trimer and hexamer were refined to a resolution of 3.04 Å, 3.12 Å, 3.34Å and 3.19 Å, respectively (Table S1).

**Figure 1.**
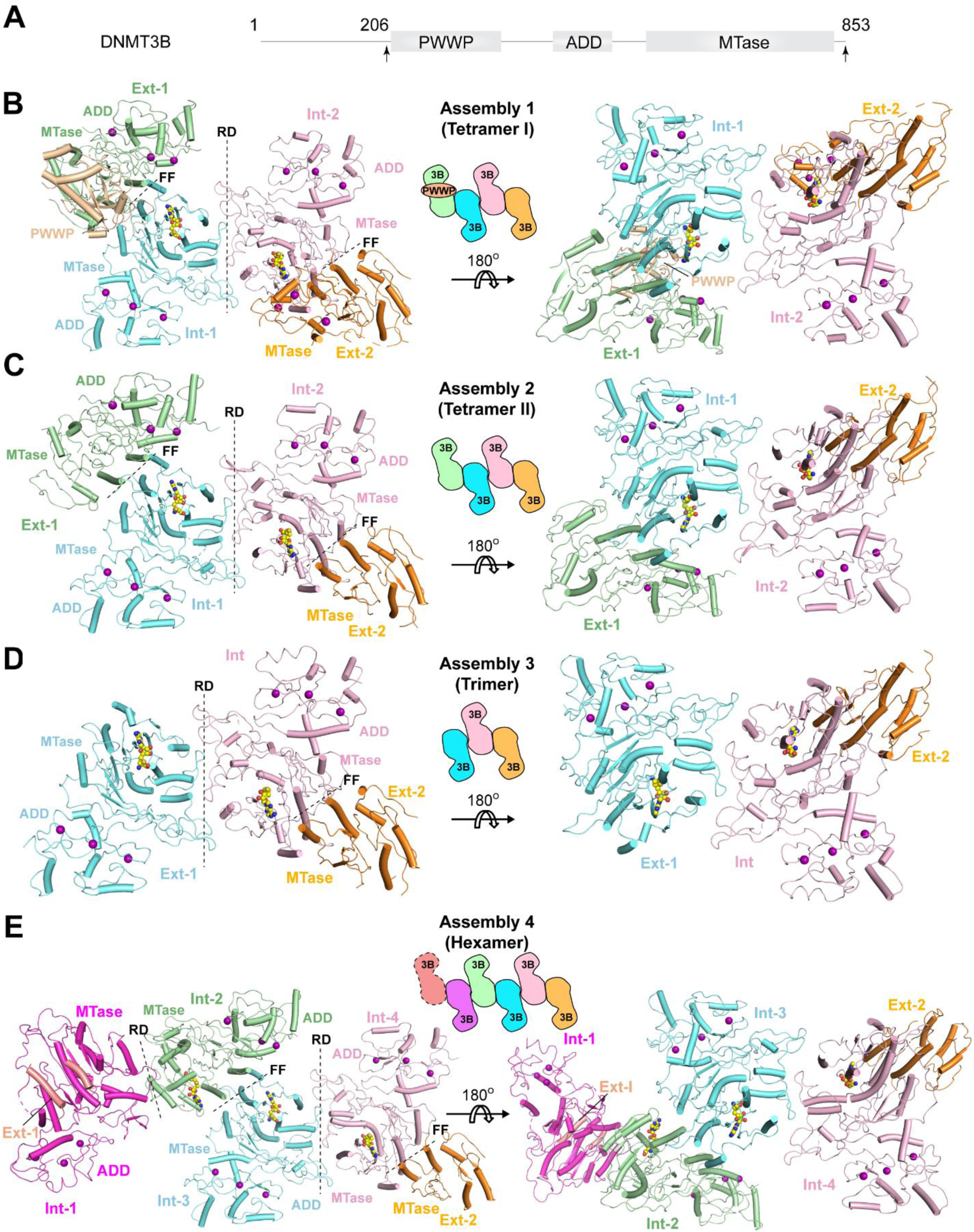
Structural overview of DNMT3B assemblies. (A) Domain architecture of human DNMT3B. The fragment used for cryo-EM structural study is indicated by arrows. (B-D) Ribbon diagram of DNMT3B tetramer I (B), tetramer II (C), trimer (D) and hexamer (E) with individual subunits color coded. Ext-1: external monomer 1, Int-1: internal monomer 1, Int-2: internal monomer 2, Int-3: internal monomer 3, Int-4: internal monomer 4, Ext-2: external monomer 2. The RD and FF interfaces are marked by dashed lines. The SAH molecules are shown in sphere representation. For clarity, schematic views of DNMT3B assemblies are shown in (B-E).

Formation of the two tetrameric forms of DNMT3B is each mediated by one RD interface and two FF interfaces, with the RD interface bridging the two central monomers and the FF interface bridging one central monomer with one external monomer (Fig. 1B,C). Such a topology of DNMT3B tetramers resembles that of the DNMT3B-DNMT3L (31,32) and DNMT3A-DNMT3L heterotetramers (16,33,39), in which the RD interface mediates the formation of DNMT3B or DNMT3A homodimer and the FF interfaces mediates the association of each DNMT3B/DNMT3A monomer with one DNMT3L molecule. Surprisingly, unlike the DNMT3B-DNMT3L heterotetramer that assumes a two-fold symmetry, the DNMT3B homotetramer is asymmetric, reflected by conformational distinctions between the two external subunits (Fig. 1B,C). The major structural difference between the two tetrameric forms lies in one of the external subunits (Ext-I), which associates with a well-defined PWWP domain in tetramer I (Fig. 1B) but not in tetramer II (Fig. 1C). However, due to the lack of density connecting this PWWP domain with any of the four ADD domains in the tetramer I, which of the tetramer I subunits gives rise to this PWWP domain is unclear. Nevertheless, based on the distances between the C-terminus of this PWWP domain and the N-termini of the ADD domains (see the details below), it is likely that the PWWP domain belongs to the central monomer next to the Ext-1 subunit of the tetramer I. Formation of the trimeric form of DNMT3B is mediated by one RD interface and one FF interface (Fig. 1D), with the three monomers packed in a similar fashion as the corresponding units of the tetramers. Formation of the hexamer is mediated by two RD interfaces and three FF interfaces. The conformation of the hexamer resembles that of tetramer II, except for the presence of a fifth subunit (Int-1) that associates via the RD interface, and residual density of a sixth subunit (Ext-1) that is further attached to the fifth subunit via the FF interface (Fig. 1E).

### Cryo-EM structure of DNMT3B tetramer I

The cryo-EM density of DNMT3B tetramer I reveals a pair of DNMT3B subunits (Int-1 and Int-2), each harboring an endogenous cofactor byproduct S-adenosyl-L-homocysteine (SAH) molecule, dimerizing via the RD interface, each of which is further flanked by a SAH-lacking external DNMT3B subunit (Ext-1 or Ext-2) via the FF interface (Fig. 1B and Fig. S3-5). The lack of SAH molecule for Ext-1 and Ext-2 is presumably attributed to the structural disordering of the RD interface that led to disruption of the SAM-binding site in these subunits (Fig. 1B and Fig. S3D-G). The MTase domains of all four subunits and the ADD domains of the central subunits were refined at a resolution of ∼3 Å, while the ADD domains of the two external subunits were refined to a resolution of ∼4 Å (Fig. S3I). Each of the DNMT3B subunits adopts a compact domain conformation, in which the ADD domain, followed by a 20-amino acid long ADD-MTase linker (residues 554-573) that runs along one end of the central β-sheet of the MTase domain, stacks against the MTase domain (Fig. 1B and Fig. S4A,B). On one face of the tetramer I, the central dimer forms a continuous DNA-binding surface (Fig. 2A), similar to what was previously observed for the DNMT3B-DNMT3L complex (31,32). In fact, structural alignment of the tetramer I with the DNMT3B-DNMT3L-DNA complex (PDB 6U8P) reveals high similarity between the central subunits, with a root-mean-square deviation (RMSD) of 1.4 Å over 553 Cα atoms (Fig. 2A). However, it is also apparent that the ADD-MTase association of the DNMT3B tetramer I creates a potential clash with the DNA substrates (Fig. 2A), suggesting that the tetramer I represents an autoinhibitory state of DNMT3B.

**Figure 2.**
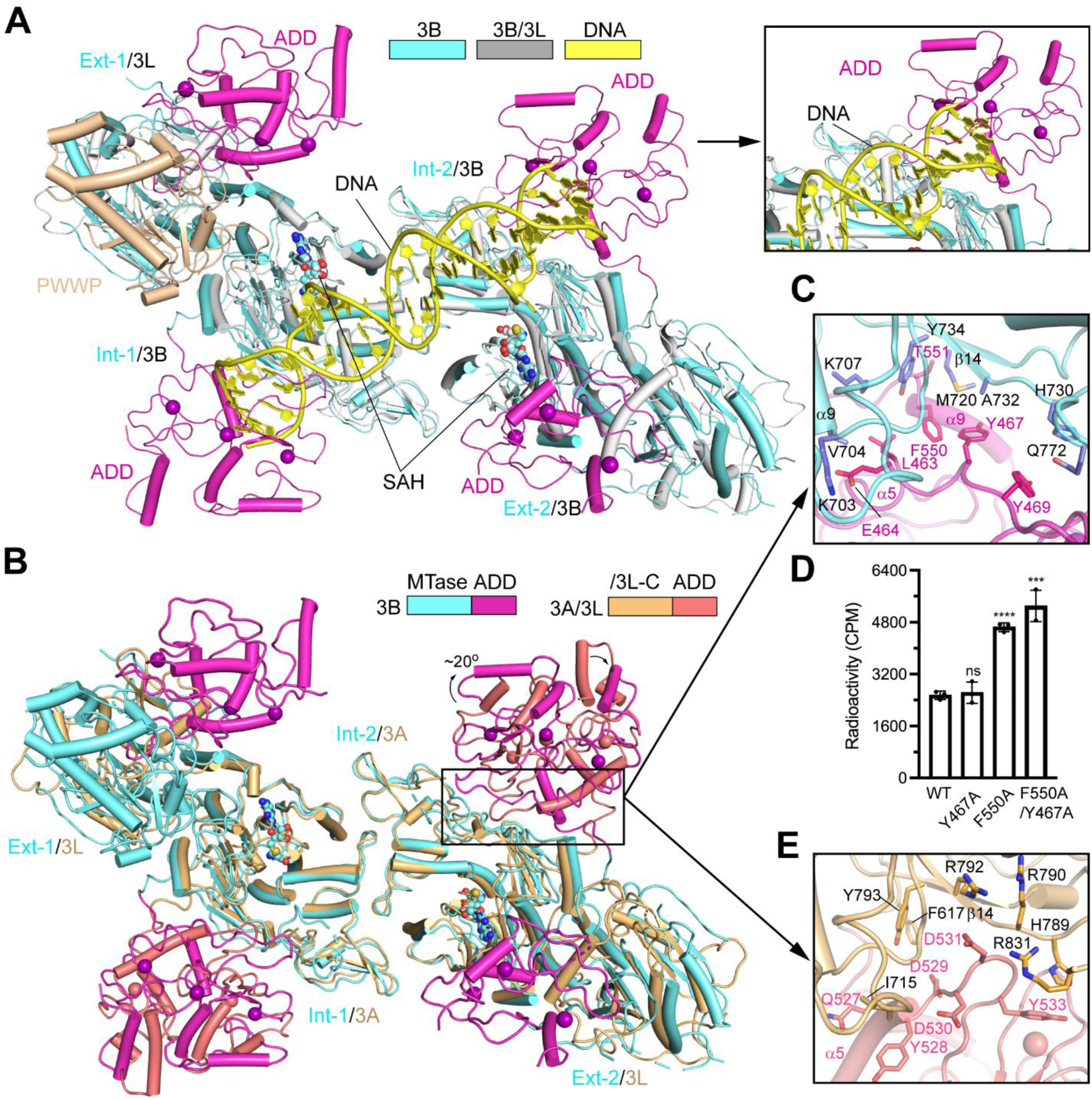
The structure of DNMT3B tetramer I reveals an autoinhibitory conformation. (A) Structural alignment between DNMT3B tetramer I (aquamarine) and the DNMT3B (grey)-DNMT3L (grey)-DNA (yellow) complex (PDB 6U8P), with the steric clash between the DNA and the ADD domain of the Int-2 subunit highlighted in the expanded view. The SAH molecules with elements color coded and zinc ions (purple) are shown in sphere representation. (B) Structural alignment between DNMT3B tetramer I (aquamarine) and the DNMT3A-DNMT3L (yellow orange) complex (PDB 4U7P). The ADD domains for DNMT3B and DNMT3A are colored in magenta and salmon, respectively. (C) Close-up view of the DNMT3B ADD-MTase interaction. (D) *In vitro* DNA methylation assay of wild-type (WT) or mutant DNMT3B PWWP-ADD-MTase fragment. Data are mean ± s.d. (n = 3 biological replicates). The two-tailed Student t-tests were performed to compare the DNMT3B WT vs mutant. ns, not significant; ***, p<0.001; ****, p<0.0001. (E) Close-up view of the DNMT3A ADD-MTase interaction. The side chains of the interaction residues in (C, E) are shown in stick representation.

Cryo-EM analysis of the tetramer I further reveals that a cluster of density, primarily extending from the N-terminus of the ADD domain of Ext-1, arches over the DNA-binding surface of DNMT3B central dimer (Fig. S4C,D). Whereas the weak intensity of this density limits it from reliable modeling, even with the 3D variability analysis by cryoSPARC (46,53), its apparent extension from the N-terminus of the ADD domain of Ext-1 suggests that this density may at least partially arise from the N-terminal region of Ext-1, encompassing the PWWP domain and/or the PWWP-ADD linker. This observation, together with the analysis the inter-domain distancing within the tetramer I (Fig. S4E), raises a possibility that the PWWP domain observed in the tetramer I may arise from the Ext-1-adjacent central subunit (Int-1). Further structural characterization is warranted to clarify the identity of this PWWP domain.

### Structural basis for the autoinhibitory state of DNMT3B

The association between the ADD and MTase domains of DNMT3B is reminiscent of what was observed for the autoinhibitory state of the DNMT3A-DNMT3L complex (39). Indeed, structural alignment of the tetramer I with DNMT3L-bound DNMT3A ADD-MTase fragment (PDB 4U7P) shows that the ADD domains of DNMT3B and DNMT3A are positioned similarly on top of their MTase domains (Fig. 2B), suggesting that the ADD domain of DNMT3B may regulate the activity of DNMT3B in a similar fashion as that of DNMT3A. Nevertheless, detailed analysis of the ADD-MTase interfaces reveals distinct domain positionings between DNMT3A and DNMT3B, with the DNMT3B ADD domain rotating away from the DNMT3A ADD domain by ∼20° (Fig. 2B). In essence, the ADD-MTase interaction within the DNMT3B tetramer I is mainly mediated by residues surrounding the α5- and α9-helices of ADD domain and the α14-helix and β14-strand of the MTase domain (Fig. 2C), involving both polar and non-polar contacts between the ADD (residues L463, E464, Y467, Y469, F550 and T551) and MTase (residues K703, V704, K707, M720, H730, A732, Y734 and Q772) domains, leading to partial occlusion of the DNA binding sites (e.g. Q772) of the MTase domain (Fig. 2C) (31). Consistent with the observations, our *in vitro* DNA methylation assays of DNMT3B PWWP-ADD-MTase indicate that introducing the ADD-MTase interface mutation F550A increased *in vitro* DNA methylation efficiency of DNMT3B by 1.8-fold (Fig. 2D and Fig. S6A,B). Although introducing another interface mutation Y467A failed to affect DNMT3B activity appreciably, introducing the Y467A/F550A double mutation led to a ∼2.0-fold activity increase of DNMT3B (Fig. 2D). The caveat of these assays is that they were based on single time point, rather than rate measurement. Consistently, our thermal shift analysis indicates that introducing the Y467A, F550A and Y467A/F550A mutations each led to reduction of melting temperature (Tm) of DNMT3B PWWP-ADD-MTase fragment by 2-3 °C (Fig. S6C,D), supporting the roles of these residues in the ADD-MTase interaction. In contrast, the ADD-MTase interaction within DNMT3A does not involve the α9-equivalent helix; rather, it is dominated by the electrostatic contacts between the potential H3-binding sites: the D529-D531 motif located C-terminal to α5-helix of the ADD domain, and the potential DNA-binding sites: residues R790 and R792 from the β14-strand of the MTase domain (Fig. 2E and Fig. S7) (39). In addition, the ADD-MTase association of DNMT3A is supported by van der Waals contacts involving residues Q527, Y528 and Y533 of the ADD domain and residues H789 and R831 of the MTase domain (Fig. 2E).

Sequence comparison of DNMT3B and DNMT3A revealed that the ADD-MTase interface residues are highly conserved between DNMT3B and DNMT3A (Fig. S7). We therefore further compared the structure of DNMT3B tetramer I with that of previously reported DNMT3A2-DNMT3B3-nucleosome complex (PDB 6PA7) (54). The DNMT3A2 and DNMT3B3 subunits of the DNMT3A2-DNMT3B3-nucleosome complex are aligned well with the internal and external subunits of the tetramer I, respectively, with the TRD-lacking DNMT3B3 resembling the external subunits more than the internal subunits of the tetramer I (Fig. S8A-C). Although the regulatory domains of DNMT3A2-DNMT3B3 were not modeled in the DNMT3A2-DNMT3B3-nucleosome complex, docking the tetramer I into the cryo-EM density of the DNMT3A2-DNMT3B3-nucleosome complex (EMD-20281) reveals a similar ADD-MTase domain conformation, in which the ADD domain is positioned in proximity with both the MTase domain and the histone H3 tail, allowing for a potential coupling between H3K4me0 readout and DNMT3A or DNMT3B activation (Fig. S8D). Interestingly, compared with the DNMT3A ADD domain in DNMT3A-DNMT3L complex (PDB 4U7P), the ADD domain of DNMT3B tetramer I appears to fit better with the cryo-EM density of the DNMT3A2-DNMT3B3-nucleosome complex (Fig. S8D-F), suggesting a closer resemblance of the ADD conformation between the DNMT3A2-DNMT3B3 complex and DNMT3B tetramer I. The molecular basis for the subtle distinction of the ADD domain conformation between DNMT3B tetramer I, DNMT3A-DNMT3L and DNMT3A2-DNMT3B3 is yet to be characterized.

### A role of the PWWP domain in the allosteric regulation of DNMT3B

Uniquely observed in tetramer I, the PWWP domain is positioned at the interface between the ADD and MTase domains of one external subunit (Ext-1), engaging in domain interactions with both the ADD and MTase domains (Fig. 3A). The PWWP-ADD association is mediated by residues K234, S273 and K276 of the PWWP domain and residues D470, D472 and Y474 of the ADD domain, involving hydrogen-bonding, electrostatic and van der Waals contacts (Fig. 3A and Fig. S4F). Note that these interactions are located immediately next to residue S270, mutation of which into proline (S270P) has been identified in patients with ICF syndrome (55). The PWWP-MTase association is mediated by the van der Waals contacts between residues M255 and M258 of the PWWP domain and catalytic-loop residues N658 and R661 of the MTase domain (Fig. 3A). Meanwhile, residue R661 forms hydrogen bonds with residues D470 and Y477 of the ADD domain, which further supports the PWWP-ADD-MTase association. The functional consequence of the PWWP-ADD-MTase association remains to be characterized. Nevertheless, structural comparison of the Ext-1 subunit of tetramer I with the previously characterized DNMT3B-DNMT3L-DNA complex reveals that the PWWP domain, like the ADD domain, would clash with the DNA substrate (Fig. 3B). Furthermore, the PWWP-ADD-MTase interaction results in a helical conformation of catalytic-loop residues K662-Y665 of Ext-1 that are occluded from DNA binding (Fig. 3A,B and Fig. S6A). Together, these observations suggest that the PWWP domain may further stabilize the autoinhibitory conformation of the external subunits of tetramer I.

**Figure 3.**
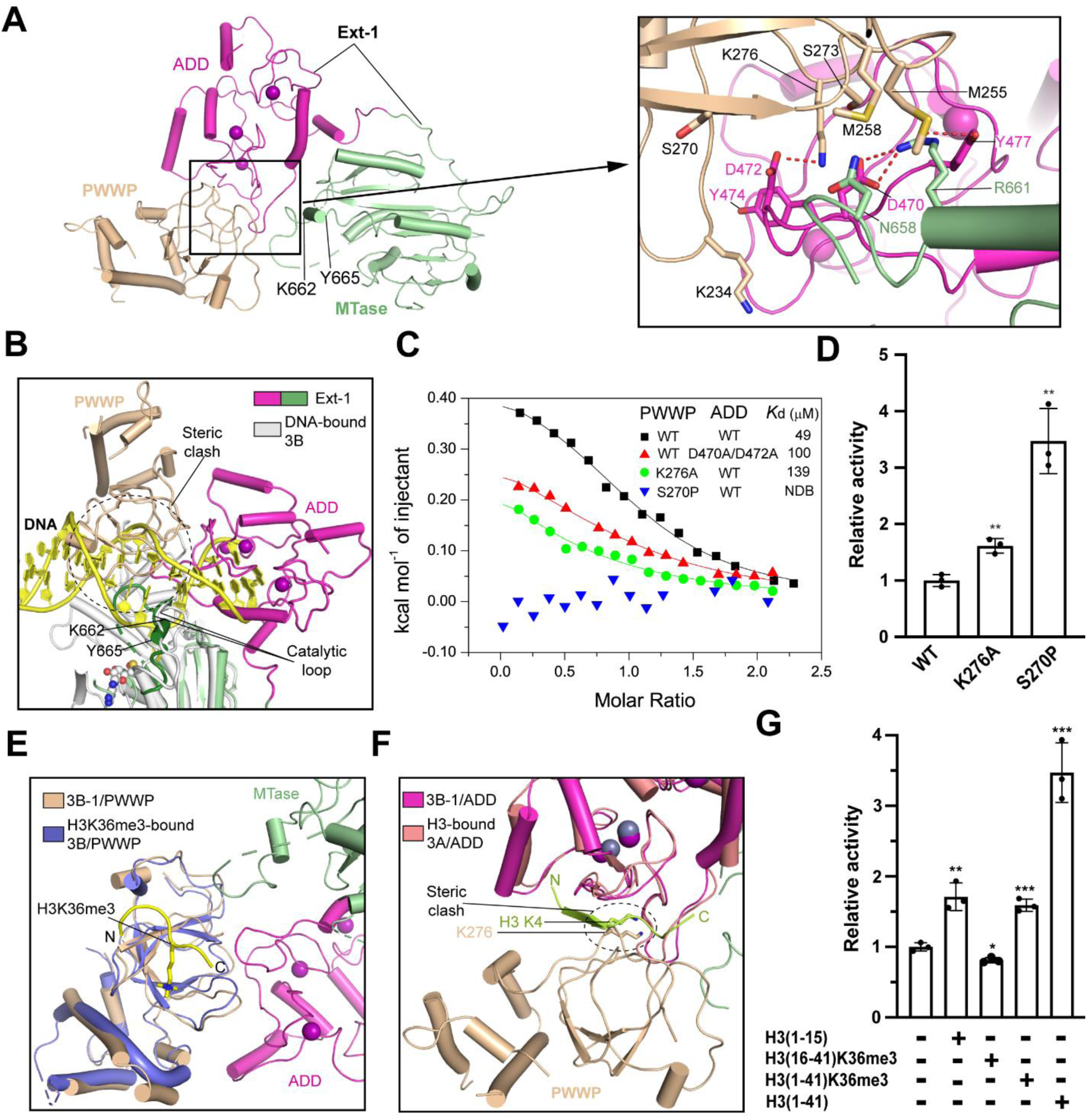
The PWWP-ADD-MTase interaction within the Ext-1 subunit of the tetramer I. (A) Ribbon representation of the association between the PWWP domain (wheat) and the ADD (magenta) and MTase (light green) domains of the Ext-1 subunit, with the detailed interactions shown in the expanded view. The hydrogen bonds are shown as dashed lines. (B) Structural alignment between DNA-bound DNMT3B MTase domain (grey, PDB 6U8P) and the PWWP-bound Ext-1 subunit. The steric clash between the PWWP domain and the DNA is indicted by dashed circle. Note that the helical conformation assumed by residues K662-Y665 on the catalytic loop (green) of Ext-1 is distinct from the corresponding region (grey) of the DNA-bound DNMT3B. The disordered segment of the catalytic loop of Ext-1 is shown as a dashed line. (C) ITC binding curves for the PWWP and ADD domains, WT or mutant. Mean and average were derived from two independent measurements. NDB, no detectable binding. (D) *In vitro* DNA methylation activities of WT or mutant DNMT3B (206-853). Data are mean ± s.d. (n = 3 biological replicates). The two-tailed Student t-tests were performed to compare WT vs mutant. **, p<0.01. (E) Structural alignment between the PWWP domain of the tetramer I and the H3K36me3 peptide (yellow)-bound PWWP domain of DNMT3B (PDB 5CIU). (F) Structural alignment between the Ext-1 ADD domain of the tetramer I and the H3K4me0 peptide (limon)-bound ADD domain of DNMT3A (PDB 3A1B). The steric clash between residues H3 K4 and PWWP K276 are indicated by dashed circle. (G) *In vitro* DNA methylation activities of DNMT3B (206-853) free and in the presence of various histone peptides. Data are mean ± s.d. (n = 3 biological replicates). The two-tailed Student t-tests were performed to compare peptide-free vs peptide-present. *, p<0.05; **, p<0.01; ***, p<0.001.

The observed PWWP-ADD-MTase association raises a possibility that the PWWP domain may contribute to the allosteric regulation of DNMT3B external and likely also internal subunits. To test this possibility, we purified the PWWP and ADD domains separately and evaluated their interaction by Isothermal Titration Calorimetry (ITC). Consistent with the structural observation, wild-type (WT) PWWP and ADD domains interacted with a dissociation constant (*K*d) of 49 ± 4 µM (Fig. 3C and Table S2). In contrast, introducing the K276A mutation to the PWWP domain and the D470A/D472A double mutation to the ADD domain reduced the binding affinity by ∼3- and 2-fold, respectively, under the experimental condition (Fig. 3C). Strikingly, introducing the ICF-associated S270P mutation to the PWWP domain completely abolished the PWWP-ADD interaction (Fig. 3C), presumably owing to a structural perturbation effect of this mutation on DNMT3B as shown by our thermal shift analysis (Fig. S9A-C) and a recent study (56). Collectively, these data confirm the interaction between the PWWP and ADD domains. Furthermore, we measured the *in vitro* DNA methylation activity of DNMT3B206-853, either WT or PWWP-mutated. Relative to WT DNMT3B206-853, the K276A and S270P mutants led to 1.6- and 3.5-fold increase in DNA methylation efficiency, respectively (Fig. 3D and Fig. S9A), suggesting that the PWWP-ADD-MTase association within the external monomers may reinforce the ADD domain-mediated DNMT3B autoinhibition. The modest impact of the K276A mutation on DNMT3B activity likely reflects the fact that this mutation led to ∼3-fold reduction of the bilateral interaction between the PWWP and ADD domains, rather than the PWWP-ADD-MTase trilateral interaction. Nevertheless, these data uncover a role for the PWWP domain in the allosteric regulation of DNMT3B.

### PWWP-H3K36m3 and ADD-H3K4me0 readouts lead to differential impacts on DNMT3B activity

The PWWP and ADD domains of DNMT3A/DNMT3B have been identified as readers for H3K36me2/3 (40,42) and H3K4me0 (37) marks, respectively. To illustrate how the PWWP-ADD domain interaction impacts their respective histone-binding activities, we compared the structures of individual PWWP and ADD domains of DNMT3B with the reported complexes of DNMT3B PWWP-H3K36me3 peptide (PDB 5CIU) and DNMT3A ADD-H3K4me0 peptide (PDB 3A1B), respectively. Structural alignment of the PWWP domain with the DNMT3B PWWP-H3K36me3 complex indicates that the PWWP-H3K36me3 interaction at the histone peptide level does not compete against the PWWP-ADD binding (Fig. 3E), suggesting that the H3K36me3 binding does not impact the PWWP-ADD association within tetramer I. On the other hand, structural alignment of the ADD domain with the ADD-H3K4me0 complex indicates that the H3K4me0-binding site of the ADD domain is shielded by both the PWWP domain and the MTase domain within tetramer I (Fig. 3F), suggesting that the interaction of the ADD domain and H3 would lead to a repositioning of the ADD domain from the MTase and the PWWP domains. Due to the lack of defined density of the other three PWWP domains, it is unclear whether the other three PWWP domains engage in any interaction with the ADD domains is unclear.

To compare the functional impact of the ADD-H3K4me0 and PWWP-H3K36me3 readouts, we performed *in vitro* DNA methylation assay for DNMT3B PWWP-ADD-MTase in the presence of various histone peptides (Fig. 3G). Presence of the H3(1–15) peptide led to a marked increase in the DNA methylation activity of DNMT3B, confirming a role of ADD-H3K4me0 binding in allosteric activation of DNMT3B (Fig. 3G). Intriguingly, incubation with a longer H3(1–41) peptide led to additional 2-fold activity increase (Fig. 3G), suggesting that residues 16-41 of the H3 peptide may also play a role in relieving the ADD domain-mediated autoinhibition. In contrast, the presence of the H3(1–41)K36me3 or H3(16–41)K36me3 peptide led to much reduced activity than their corresponding H3K36me0 peptides (Fig. 3G), suggesting that the ADD-H3K4me0 readout, but not the PWWP-H3K36me3 readout, allosterically activates DNMT3B. It is worth noting that the H3(1–15) and H3(16–41)K36me3 peptides impacted the activity of DNMT3B harboring a H3K36me3 pocket mutation (W263A) (42) similarly as they did to the activity of WT DNMT3B (Fig. S9D,E), implying that the moderate activity decrease for DNMT3B in the presence of H3(16–41)K36me3 peptide was presumably attributed to non-specific DNA binding, rather than the PWWP-H3K36me3 readout. Together, these data reveal a differential regulatory effect of the two epigenetic marks: H3K4me0 and H3K36me3, in the DNMT3B-mediated DNA methylation. The caveat is that these experiments described above were performed at the histone peptide level, which may deviate from the actual nucleosome environment in cells.

### Coupled RD-interface integrity and the substrate-binding interface of DNMT3B

Assembly of DNMT3B tetramer I results in distinct chemical environments between the central and external subunits, with the Int-1 and Int-2 oligomerizing via both RD and FF interfaces and the Ext-1 and Ext-2 oligomerizing via the FF interface only. To understand how the integrity of the RD interface affects DNMT3B conformation, we compared the structures of the four subunits of the tetramer I (Fig. 4A). Superposition of the two central subunits reveals a nearly identical conformation (RMSD of <0.1 Å over 427 aligned Cα atoms), superposition of the two external subunits gives an RMSD of 1.2 Å over 213 aligned Cα atoms, superposition of the Ext-1 subunit with the two central subunits each gives an RMSD of 1.0 Å over 287 aligned Cα atoms, and superposition of the Ext-2 subunit with the two central subunits each gives an RMSD of 1.5 Å over 224 aligned Cα atoms.

**Figure 4.**
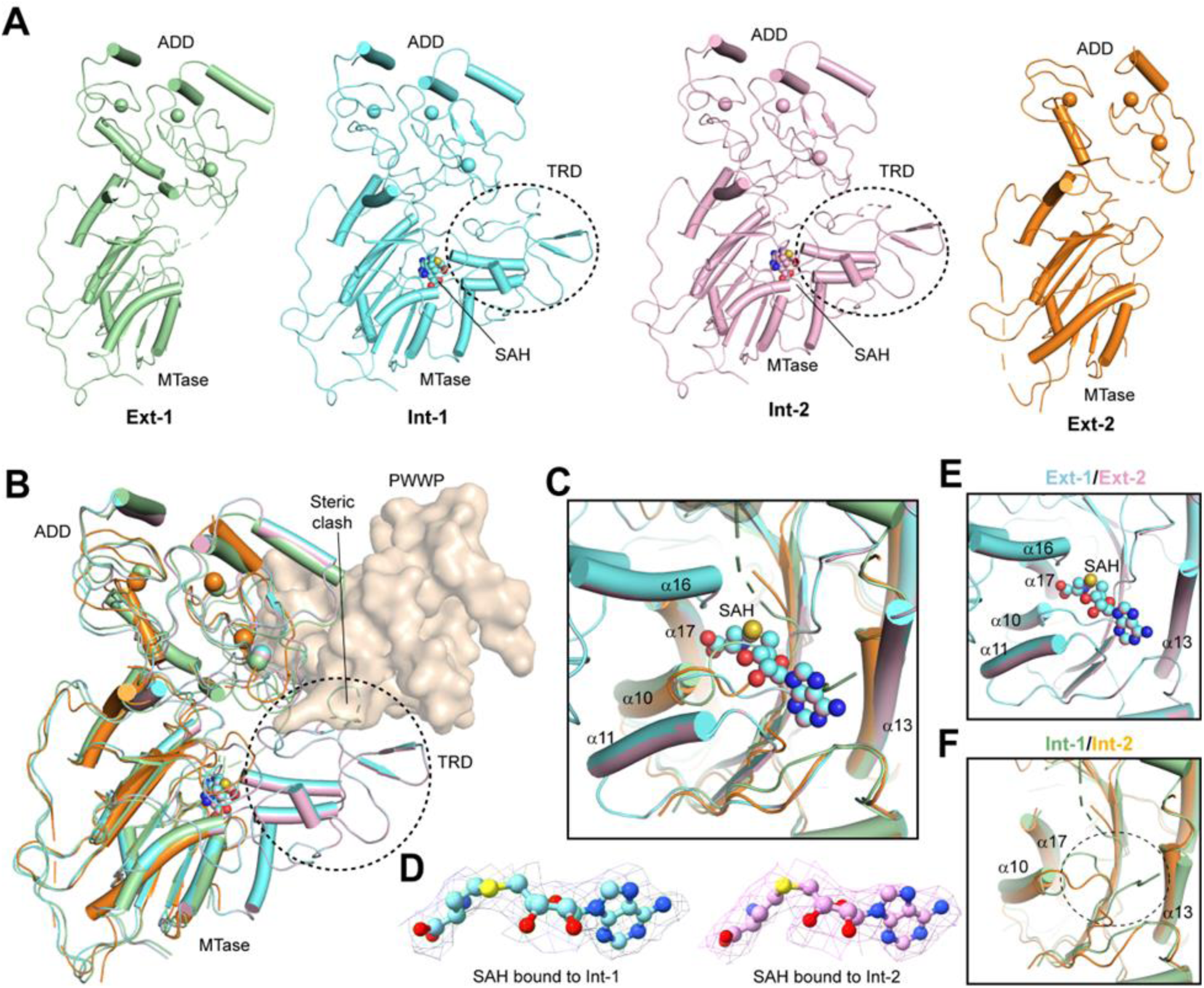
Structural comparison of the DNMT3B subunits within the tetramer I. (A) Ribbon representation of the Ext-1, Int-1, Int-2 and Ext-2 subunits of tetramer I, with the ADD and MTase domains labeled. Note that the TRD (circled with dashed line) and the SAH-binding site are folded in Int-1 and Int-2 but disordered in Ext-1 and Ext-2. (B) Structural alignment of the four subunits of the tetramer I, with the TRDs of Int-1 and Int-2 circled. The clash between the PWWP domain, shown in surface representation, and a portion of the folded TRD is indicated. (C) Close-up view of the cofactor-binding sites in the four subunits. The α-helices that are involved in the cofactor-binding pocket are labeled. (D) EM densities for the SAH molecules bound to the Int-1 and Int-2 subunits of the tetramer I. (E) Close-up view of the SAH-binding sites in the Int-1 and Int-2 subunits. (F) Close-up view of the cofactor-binding site (dashed circle) disrupted in the Ext-1 and Ext-2 subunits.

Detailed comparison of the four subunits of the tetramer I reveals different folding behaviors between the external and central subunits (Fig. 4A,B and Fig. S3C-H and Fig. S5). The density for the two central subunits is both well defined, permitting us to trace the entire ADD and MTase domains (residues 415-853) (Fig. S3E,F), except for a loop of the TRD (residues 778-787) that was shown to undergo a disorder-to-order transition upon DNA binding (Fig. S5B-D) (9,31,32). In contrast, the two external subunits lack defined density for the TRD (residues 748-801), as well as the α11-helix (residues 606-620) of the catalytic core (Fig. 4A and Fig. S3D,G), which together constitute the underlying elements of the RD interface. This observation suggests that disruption of the RD interface leads to structural disordering of the corresponding elements. Interestingly, structural alignment of the central subunits with the PWWP domain-associated Ext-I also reveals a potential steric clash between the PWWP domain and the otherwise structurally ordered TRD (Fig. 4B), suggesting of an interplay between the ADD-PWWP-MTase domain interaction and the structural disordering of the TRD of the external subunits. Consistent with this notion, our dynamic light scattering (DLS) analysis reveals that removal of the PWWP domain and/or ADD domain led to increased tendency of protein aggregation at various salt conditions (Fig. S10). Another major difference between the central and external monomers lies in the cofactor-binding pocket, formed by the loop C-terminal to the α10-helix, the α11- and α13-helices, and the α16- and α17-helices of the TRD (Fig. 4C and Fig. S5E,F). The cofactor-binding pocket is formed in each of the central subunits (Fig. 4C-E and Fig. S5E,F), reminiscent of the DNMT3B-DNMT3L complexes (31,32). In contrast, the corresponding pockets in the external subunits were disrupted, owing to the structural disordering of the α11-helix and the TRD (Fig. 4C,F and Fig. S5G,H). In addition, the loop C-terminal to the α10-helix in Ext-1 underwent a movement toward the SAH pocket, which further occludes the cofactor binding (Fig. 4C).

The lack of TRD folding of the external subunits coincides with the region that is spliced out in isoform 3 of DNMT3B (DNMT3B3) (Fig. S8A-C), which becomes enzymatically inactive yet remains capable of associating with the DNMT3A or DNMT3B homodimer into a stable tetramer for DNA methylation (Fig. S8A) (54,57).

### Conformational plasticity of the FF interface

We recently reported the crystal structure of DNMT3B MTase domain in a macro-oligomeric form, in which DNMT3B monomers stack against each other via alternative RD and FF interfaces into a left-handed helical assembly (9). Structural comparison of the tetramer I with the macro-oligomeric DNMT3B MTase (PDB 7V0E) (9) and the DNA/ DNMT3L-bound DNMT3B MTase (PDB 6U8P) (31) reveals a common theme of oligomerization that is mediated by alternating RD and FF interfaces (Fig. S11A, B). Accordingly, superposition of the central subunits of the tetramer I with the corresponding regions of the DNMT3B macro-oligomer and the DNMT3B-DNMT3L complex gives an RMSD of 1.2 Å and 1.4 Å over 509 and 542 aligned Cα atoms, respectively. Notably, all these complexes involve the α13- and α14-helices of DNMT3B for the formation of the FF interface, dominated by a hydrophobic cluster formed by residues F673, Y676, H677, Y681 and F713 (Fig. S11C-F).

Detailed comparison of DNMT3B tetramer I with macro-oligomeric DNMT3B and DNMT3B-DNMT3L further reveals distinct inter-molecular interactions underlying the FF interface. For instance, DNMT3B F673 engages in a reciprocal ring-stacking interaction in DNMT3B tetramer I and stacks with DNMT3L F261 in a similar fashion in the DNMT3B-DNMT3L complex; in contrast, it shifts to engage in sidechain packing with residue F713′ (prime symbol denotes residues from the symmetry-related subunit) in the macro-oligomeric form (Fig. S11C-F). In addition, DNMT3B residue Y676 is sandwiched by residues H667′ and Y676′ in DNMT3B tetramer I and by DNMT3L H264 and R265 in DNMT3B-DNMT3L for ring-stacking interactions; in the macro-oligomeric form of DNMT3B, it shifts toward residue Y676′ for a sidechain hydrogen-bonding interaction (Fig. S11C-F). As a result, the α13- and α14-helices of the tetramer I or DNMT3B-DNMT3L tetramer exhibit a lateral shift of 6-7 Å relative to the corresponding regions of the DNMT3B macro-oligomer (Fig. S11A, B), as well as differential buried surface areas at the FF interface (∼860 Å2 for DNMT3B tetramer I and ∼780 Å2 for DNMT3B-DNMT3L vs ∼710 Å2 for DNMT3B macro-oligomer). The functional implication of the different oligomerization states of DNMT3B and the mechanism underlying their transition awaits further investigation. It remains possible that the altered FF interface in the macro-oligomeric form of DNMT3B is attributed to a crystallization packing effect (9). Nevertheless, these observations suggest a conformational plasticity for the FF interface of DNMT3B.

### Structure of DNMT3B tetramer II

The EM density for the tetramer II permitted us to trace the ADD and MTase domains of the two central subunits (resolution of 3-4 Å) and one of the external subunits (resolution of 4-5 Å), whereas the MTase domain for the other external subunit (resolution of ∼5 Å) was only partially modeled (Fig. S12A-D). In addition, a blob of unmodeled density, which presumably arises from the N-terminal regions of DNMT3B subunits, is clustered on top of the DNA-binding groove of the central subunits (Fig. S12C).

Structural comparison of the tetramer II with the tetramer I reveals high similarity (Fig. 5A), with an RMSD of 0.6 Å over 1130 aligned Cα atoms. As with those in the tetramer I, the ADD domain of each subunit packs against the MTase domain, resulting in an autoinhibitory conformation (Fig. 5A). As mentioned earlier, the major difference between the two alternative tetramers lies in the fact that the PWWP domain is associated with one external subunit (Ext-1) in the tetramer I but becomes untraceable in the tetramer II (Fig. 5A,B). Accordingly, the catalytic loop of the Ext-1 subunit of the tetramer II lacks a helical conformation for residues K662-V665 that is otherwise present in the Ext-1 subunit of the tetramer I and becomes highly disordered (Fig. 5B). In addition, the blob of density arching over the DNA-binding surface of the central subunits appears stronger than that in the tetramer I (Fig. S12C-E), likely attributed to the repositioning of the PWWP domain. These observations suggest a degree of conformational dynamics of the PWWP domain. Similar to those in the tetramer I, the TRDs and the SAH-binding pockets of the two central subunits of the tetramer II are well folded (Fig. 5C,D and Fig. S12F), whereas the corresponding regions of the external monomers become disrupted (Fig. 5C,E). This observation reinforces the notion that the folding of the TRD and cofactor-binding pocket of DNMT3B is coupled with the integrity of the RD interface.

**Figure 5.**
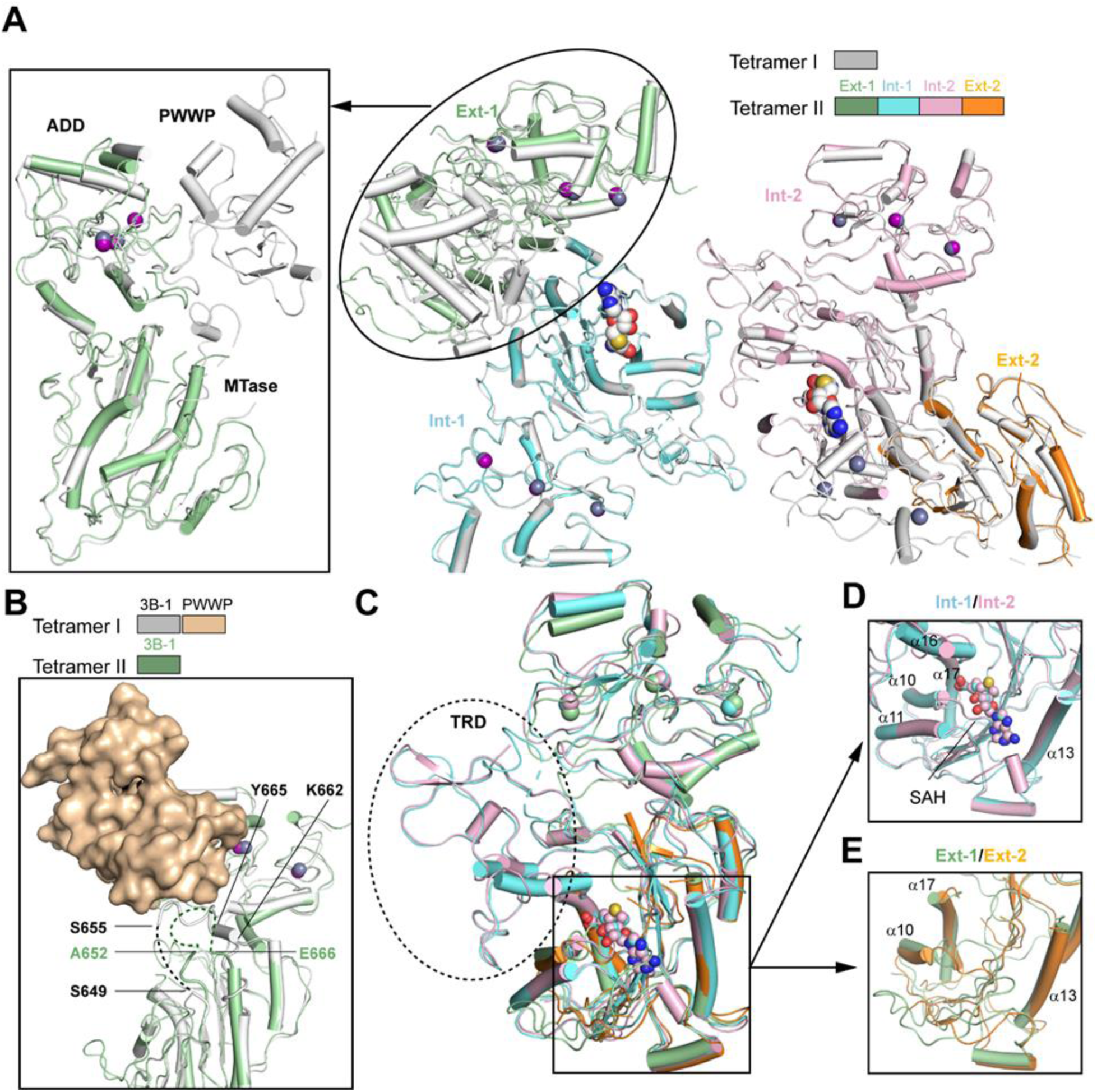
Structural analysis of DNMT3B tetramer II. (A) Structural alignment between DNMT3B tetramer II, with individual subunits color-coded, and the tetramer I (grey). The alignment between the Ext-1 subunits of the tetramer II and tetramer I is shown in the expanded view. The SAH molecules and zinc ions (purple) are shown in sphere representation. (B) Close-up view of the catalytic loops of the tetramer I and tetramer II. The disordered regions are shown as dashed lines. Note that residues K662-Y665 assumes a helical conformation in the PWWP-bound Ext-1 subunit of the tetramer I but becomes disordered in the Ext-1 subunit of the tetramer II, presumably due to a lack of PWWP domain contact. (C) Structural alignment of the four subunits of the tetramer II, with the TRD and SAH-binding sites folded in the Int-1 and Int-2 subunits are marked by dashed circle and solid rectangle, respectively. (D) Close-up view of the SAH-binding sites in the Int-1 and Int-2 subunits. (E) Close-up view of the cofactor-binding site (dashed circle) disrupted in the Ext-1 and Ext-2 subunits.

### Structural basis of the trimeric and hexameric assemblies of DNMT3B

The co-existence of DNMT3B trimer and hexamer with the tetramers suggests a dynamic equilibrium between the assembly states of DNMT3B. The DNMT3B trimer is formed by two subunits with defined ADD and PWWP domains (Ext-1 and Int) that dimerize via the RD interface, which is further joined via the FF interface by another subunit (Ext-2), in which only a portion of the MTase domain was traced (Fig. S13). The hexameric assembly reveals three subunits (Int-2 to Int-4) with defined ADD and MTase domains, flanked by two subunits that either partially (Int-1) or completely (Ext-2) lack the density of the ADD domain (Fig. S14). In addition, we were able to identify the residual density of a sixth subunit (Ext-1), corresponding to the α13 and α14-helices of the MTase domain, which are attached to the Int-1 subunit via the FF interface (Fig. S14C). The trimer shows high similarity with the corresponding assemblies within the tetramers, with an RMSD of 0.72 Å over 810 aligned Cα atoms between the trimer and the unit formed by subunits Int-1, Int-2 and Ext-2 of the tetramer I (Fig. 6A). Likewise, among the six subunits of the hexamer, subunits Int-2 to Int-5 align well with tetramer I (Fig. 6B) and II, giving an RMSD of 0.79 Å and 0.82 Å over 1112 and 1115 aligned Cα atoms, respectively.

**Figure 6.**
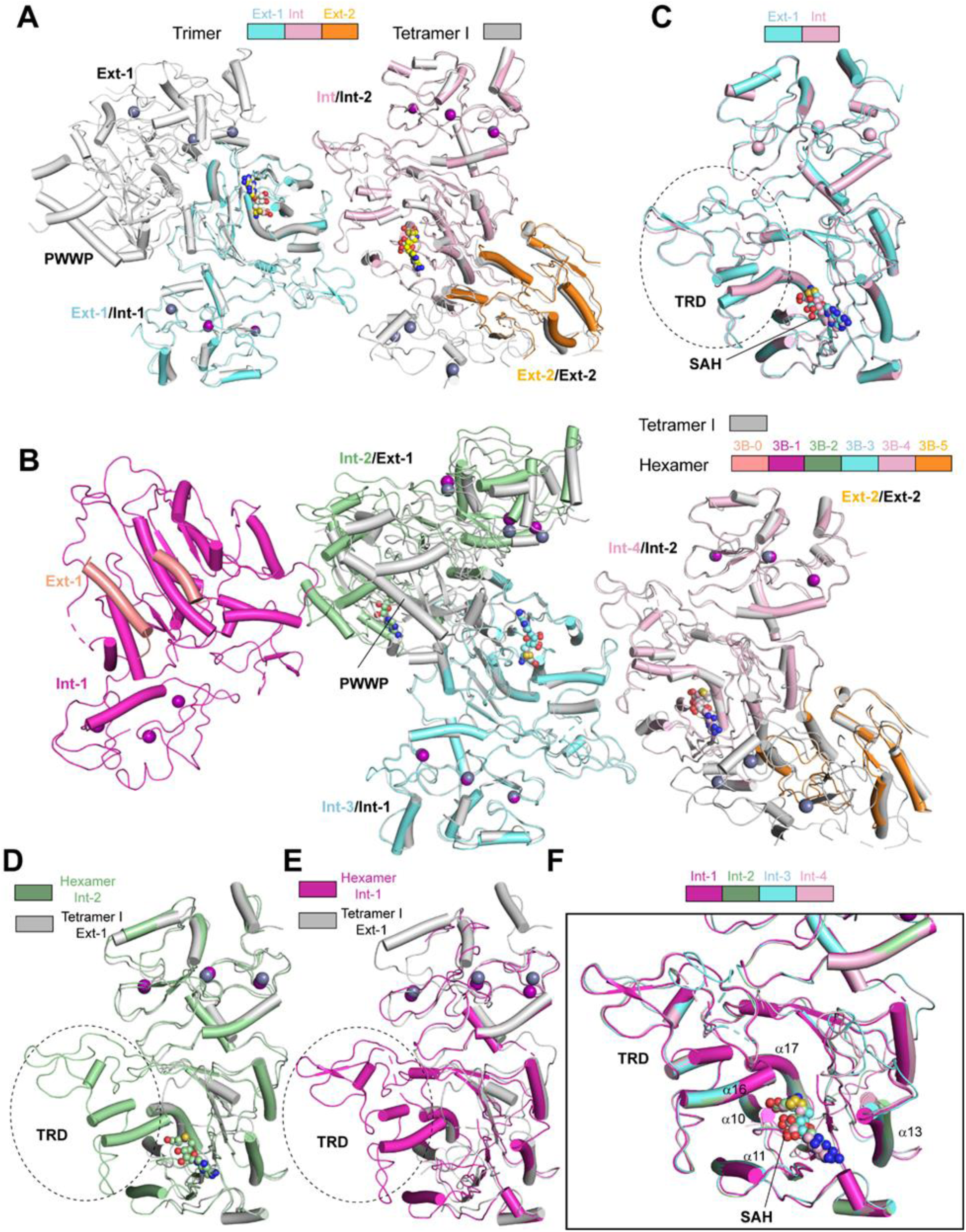
Structural analysis of DNMT3B trimer and hexamer. (A) Structural alignment between DNMT3B trimer, with individual subunits color coded, and the tetramer I (grey). (B) Structural alignment between DNMT3B hexamer, with individual subunits color coded, and the tetramer I (grey). (C) Structural alignment of the Ext-1 and Int subunits of the trimer, with the TRD and bound-SAH molecules labeled. (D-E) Structural alignment between the Int-2 (D) or Int-1 (E) subunit of the hexamer and the Ext-1 subunit of the tetramer I. Note that the TRD is folded in the Int-1 and Int-2 subunits of the hexamer, but not in the Ext-1 subunit of the tetramer I. (F) Structural alignment of the Int-1, Int-2, Int-3 and Int-4 subunits of the hexamer, highlighting the folding of the TRD and SAH-binding site in these molecules.

In the trimer, the two RD interface-related subunits (Ext-1 and Int) each contain a folded TRD and SAH-binding pocket (Fig. 6C), resulting in a largely preserved substrate-binding surface (Fig. S13E). On the other hand, the RD interface-disrupted Ext-2 subunit does not contain an intact SAH pocket, reminiscent of what was observed for the tetramers (Fig. 4C). Likewise, in the hexamer, the internal subunits Int-1, Int-2, Int-3, Int-4 and Int-5, but not the external subunits Ext-1 and Ext-2, each contain a folded TRD and cofactor-binding pocket, resulting in two discrete DNA-binding surfaces that potentially mediate binding of a pair of parallel substrates (Fig. 6C-F and Fig. S14F), as proposed previously (12). Together, these observations reinforce the notion on the coupling between the integrity of the RD interface and the folding of the TRD and the cofactor-binding pocket. Note that the PWWP domain, like that in the tetramer II and trimer, becomes untraceable in the hexamer (Fig. S14C), suggesting that the transition from the tetramer I into the tetramer II, accompanied by the repositioning of the PWWP domain, is likely the first step for the transition from the tetramer I to a higher-order DNMT3B oligomer (e.g hexamer). It is also worth mentioning that a blob of untraceable density, which presumably arises from the N-terminal domains of the subunits, is also clustered on top of the DNA-binding groove of the trimer and hexamer (Fig. S13C and Fig. S14C), suggesting the association between the N-terminal region of DNMT3B and its DNA binding grooves as a recurrent mechanism among the DNMT3B assemblies.

Structural comparison of the FF interface-disrupting Ext-1 subunit of the trimer with the FF interface-engaging Int and Ext-2 subunits reveals a similar conformation at the FF interface-surrounding region (e.g. α13 and α14-helices), suggesting that dissociation of the tetramers into trimer via the FF interface disruption does not lead to TRD unfolding of the newly exposed external subunit (Ext-2). Whether the trimer is in part stabilized by chemical crosslinking treatment during sample preparation and whether it remains catalytically active awaits further investigation. Nevertheless, as with that in the tetramers, the catalytic loops of the external subunits appear consistently more disordered than those of the internal subunits in the trimer, the tetramers and the hexamer (Fig. S6A and Fig. S15A-C), suggesting that both RD and FF interface impact the conformational dynamics of DNMT3B, albeit to a different extent.

## Discussion

Oligomeric assembly provides an important mechanism in regulating DNMT3A- and DNMT3B-mediated DNA methylation (34,35). While the heterotetrameric complex of DNMT3A and DNMT3B with DNMT3L has been well elucidated, the underlying mechanism for homo-oligomerization of DNMT3A and DNMT3B remains elusive. Furthermore, currently reported structural characterizations of the DNMT3A and DNMT3B complexes mainly focus on their C-terminal MTase domains. How the N terminal domains cooperate with the MTase domain in controlling the assembly and DNA methylation activities of DNMT3A and DNMT3B is poorly understood. This study, through structural and biochemical characterization of DNMT3B under various assembly states, provides key insights into the allosteric regulation and the dynamic assembly of DNMT3B homo-oligomers.

First, the cryo-EM structure of a DNMT3B fragment spanning the PWWP, ADD and MTase domains provides a framework for understanding the functional regulation of DNMT3B by its N-terminal domains. Importantly, the N-terminal ADD domain of DNMT3B closely packs against its C-terminal MTase domain, resulting in partial occlusion of the DNA-binding sites of the MTase domain. The ADD-MTase domain interaction of DNMT3B resembles what was previously observed for the autoinhibitory DNMT3A-DNMT3L complex (39), suggesting that DNMT3B homo-oligomer may adopt a similar autoinhibitory mechanism as that of the DNMT3A-DNMT3L complex. On the other hand, it is also apparent that DNMT3B engages in a distinct set of residues for the ADD-MTase interaction than DNMT3A in DNMT3A-DNMT3L: in contrast to the ADD domain of DNMT3A in DNMT3A-DNMT3L that engages the H3-binding sites (e.g. residues D528 and D530) for an interaction with the MTase domain, the ADD domain of DNMT3B presents the loop residues preceding the H3-binding sites for the interaction with the MTase domain. It remains possible that the ADD-MTase interface of DNMT3B may partially overlap with H3-binding sites in the context of the PWWP-ADD-MTase fragment, given that our *in vitro* DNA methylation assay indicated that residues beyond H3 (1–15) also play a role in allosteric activation of DNMT3B. Furthermore, our structural modeling analysis of DNMT3B tetramer I in complex with nucleosome, using the DNMT3A2-DNMT3B3-nucleosome complex as a template, reveals that DNMT3B ADD domain, as with DNMT3A ADD domain, is positioned near to H3 tail when bound to nucleosome, permitting a coupling between H3K4me0 readout and allosteric activation of DNMT3B. Interestingly, the conformation of DNMT3A ADD in the context of the DNMT3A2-DNMT3B3 complex appears more similar to that of DNMT3B tetramer I than that of DNMT3A in the context of DNMT3A-DNMT3L complex, supporting the functional relevance of the ADD conformation in this study. Whether the distinct ADD-MTase domain interaction between DNMT3B and DNMT3A-DNMT3L is attributed to distinct conformational states remains to be determined.

Second, this study provides additional functional insight into the PWWP domain. Through combined structural and biochemical analysis, this study unexpectedly identified that the ADD and MTase domains of one external subunit of the DNMT3B tetramer I are associated with one PWWP domain. This inter-domain interaction leads to occlusion of the H3K4me0-binding sites of the ADD domain but not the H3K36me3-binding sites of the PWWP domain, which might impact chromatin targeting of DNMT3B. Along the line, introducing the DNMT3B K276E mutation at the PWWP-ADD interface was recently shown to increase the heterochromatin localization of DNMT3B (56), supporting a link between the PWWP-ADD interaction and the targeting specificity of DNMT3B. Furthermore, the PWWP-ADD-MTase interaction appears to help stabilize DNMT3B homotetramer, as indicated by our structural and DLS analysis, thereby influencing the dynamic equilibrium of DNMT3B assembly. In addition, the PWWP domain, through interaction with both the ADD domain and the catalytic loop of DNMT3B, might reinforce the ADD domain-mediated autoinhibition. In these contexts, previous studies have demonstrated that the ICF mutation S270P, located near to the PWWP-ADD interface, impairs the interaction between DNMT3B and H3K36me3-marked nucleosome *in vitro* (44), reduces the stability of DNMT3B in cells (58), and affects the recruitment of DNMT3B to the major satellite repeats at pericentric heterochromatin (59). This study showed that this mutation causes abolished PWWP-ADD interaction and reduced thermal stability of DNMT3B. Whereas the former may boost the DNA methylation activity of DNMT3B *in vitro*, the latter presumably leads to decreased DNMT3B level and DNA hypomethylation in cells (55,56). It is worth mentioning that out of the four PWWP domains in the tetramer I, only one PWWP domain was traced. Whether the other three PWWP domains engage in transient inter-domain interactions with the ADD and/or MTase domains in tetramer I awaits further investigation.

Our structural study also reveals a blob of untraceable density clustered on top of the DNA-binding surface of all DNMT3B assemblies, which may contribute to the asymmetric nature of the DNMT3B structures. The notion that this density belongs to the N-terminal region of DNMT3B, i.e. the PWWP domain and the PWWP-ADD linker, is supported by the observation that it becomes more evident in the DNMT3B tetramer II, trimer and hexamer, which lack a defined PWWP domain. Given the proximity of this cluster of density with the DNA-binding surface and its interplay with the PWWP domain, it is conceivable that this density may affect both the DNA binding and oligomerization of DNMT3B, thereby contributing to another layer of DNMT3B regulation. Further biochemical and cellular assays are warranted to decipher the functional regulation by the N-terminal domains of DNMT3B.

Finally, this study provides the mechanistic basis for the dynamic assembly of DNMT3B, uncovering a coupling between DNMT3B oligomerization and its substrate binding. Our previous studies revealed a disorder-to-order transition for the TRD loop upon DNA binding, and the differential interaction between the TRD loop and the DNMT3A R882-equivalent residues on the RD interface underpins the distinct CpG recognition modes between DNMT3A and DNMT3B (31,33). Here, structural comparison of these complexes reveals a previously unappreciated interplay between the RD interface, TRD and the co-factor binding pocket: the subunits with a disrupted RD interface are associated with disordered TRD and cofactor-binding pocket, while continuing oligomerization via the RD interface, i.e. transition from the tetramers to the hexamer, leads to restoration of their structural ordering. This RD interface-coupled folding of the TRD and the cofactor-binding pocket of DNMT3B provides a mechanism in restricting the activities of the external subunits of the tetrameric assemblies, which may help regulate substrate and cofactor bindings of DNMT3B. Along the line, the lack a RD-interface-engaging TRD in the MTase-like domains of DNMT3L and DNMT3B3 leads to their dominant tetrameric assembly with DNMT3A/DNMT3B (16,19,54,57), which may provide a mechanism for regulating the activity, stability and target specificity of the latter for tissue- and development stage-specific DNA methylation (13–15,18,45,57,60).

Previous studies have indicated that the oligomerization states of DNMT3A and DNMT3B need to be well balanced: disruption of the DNMT3A or DNMT3B tetramer led to loss of their DNA methylation activities *in vitro* and in cells (9–12,16,17), whereas macro-oligomerization promoted by the DNMT3A R882 mutations causes aberrant chromatin targeting and enzymatic activity of DNMT3A (22–24,30,61–65), which is associated with pathogenesis of acute myeloid leukemia (21). In this regard, this study provides a framework for future therapeutic intervention of DNMT3A/DNMT3B-associated diseases.

## Data availability

The 3D cryo-EM maps have been deposited in the Electron Microscopy Data Bank under the accession numbers EMD-28156, EMD-28157, EMD-28158 and EMD-28159. Atomic coordinates for the structural models have been deposited in the Protein Data Bank under accession code 8EIH, 8EII, 8EIJ and 8EIK.

## Acknowledgments

We thank staff members at the Stanford-SLAC Cryo-EM Center (S2C2), which is supported by the National Institutes of Health Common Fund Transformative High-Resolution Cryo-Electron Microscopy program (U24 GM129541), for cryo-EM data collection.

## Author Contributions

J.L performed experiments and data analysis. J.F. contributed to biochemical analysis. H.Z. assisted in data processing. K.L. and N.K. assisted in protein purification. J.S. conceived the project and performed data processing and analysis. J.L. and J.S. prepared the manuscript.

