## Supplementary Information for "Structural basis for the allosteric regulation and dynamic assembly of DNMT3B"

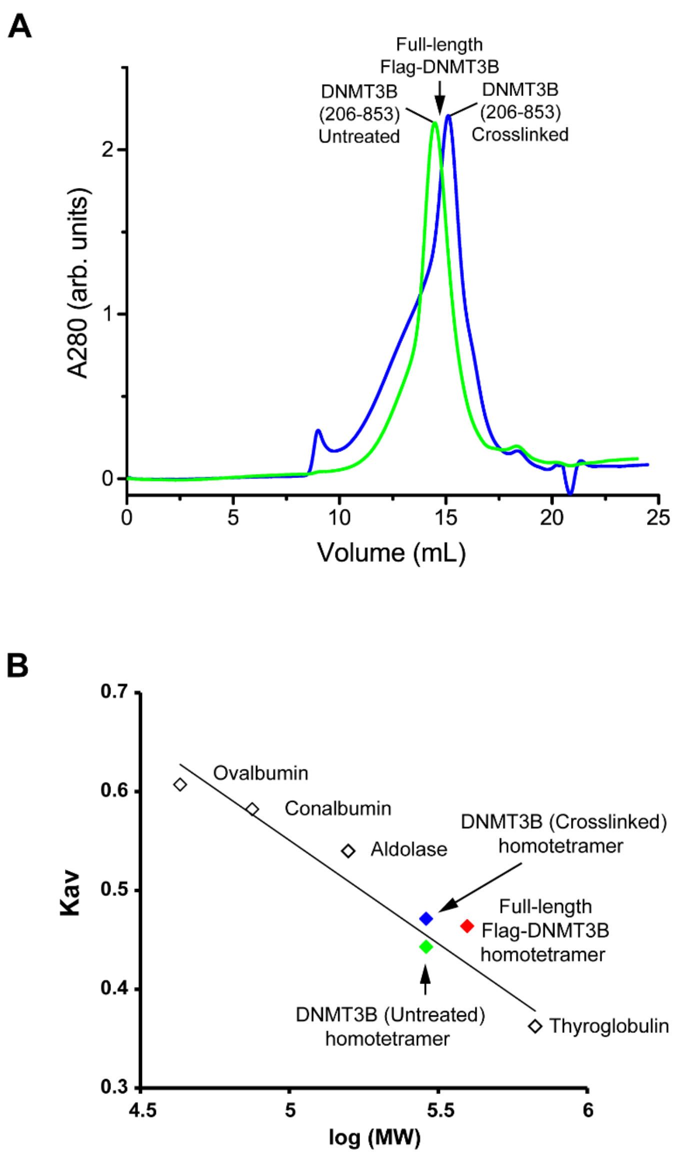


**Figure S1. Size-exclusion chromatography analysis of DNMT3B (206-853) protein.** (A) Elution profile of DNTM3B (206-853), either untreated or crosslinked with 0.02% Glutaraldehyde, on a Superose 6 increase 10/300 column. The elution volume for full-length Flag-tagged DNMT3B purified from mouse embryonic stem cells, as determined previously (*1*), is marked by arrow. (B) The Superose 6 column, as previously calibrated with Thyroglobulin, Aldolase, Conalbumin, and Ovalbumin proteins (*1*), was used to estimate the oligomeric state of DNMT3B (206-853). As previously described (*1*), the calibration curve was obtained by fitting the gel phase distribution coefficient (Kav) with the logarithm of molecular weight, log (MW). The elution volume of each protein was used to calculate the Kav using the formula: Kav = (Ve - Vo)/(Vc – Vo), where Ve = elution volume of protein, Vo = column void volume, Vc = geomtric column volume. The Thyroglobulin, Aldolase, Conalbumin, and Ovalbumin proteins are marked as open diamonds in accordance with their respective coordinates of Kav and log (MW). The predicted coordinates for the tetrameric forms of untreated full-length Flag-tagged DNMT3B, untreated DNMT3B (206-853), and chemically crosslinked DNMT3B (206-853) are indicated by red, green and blue diamonds, respectively.

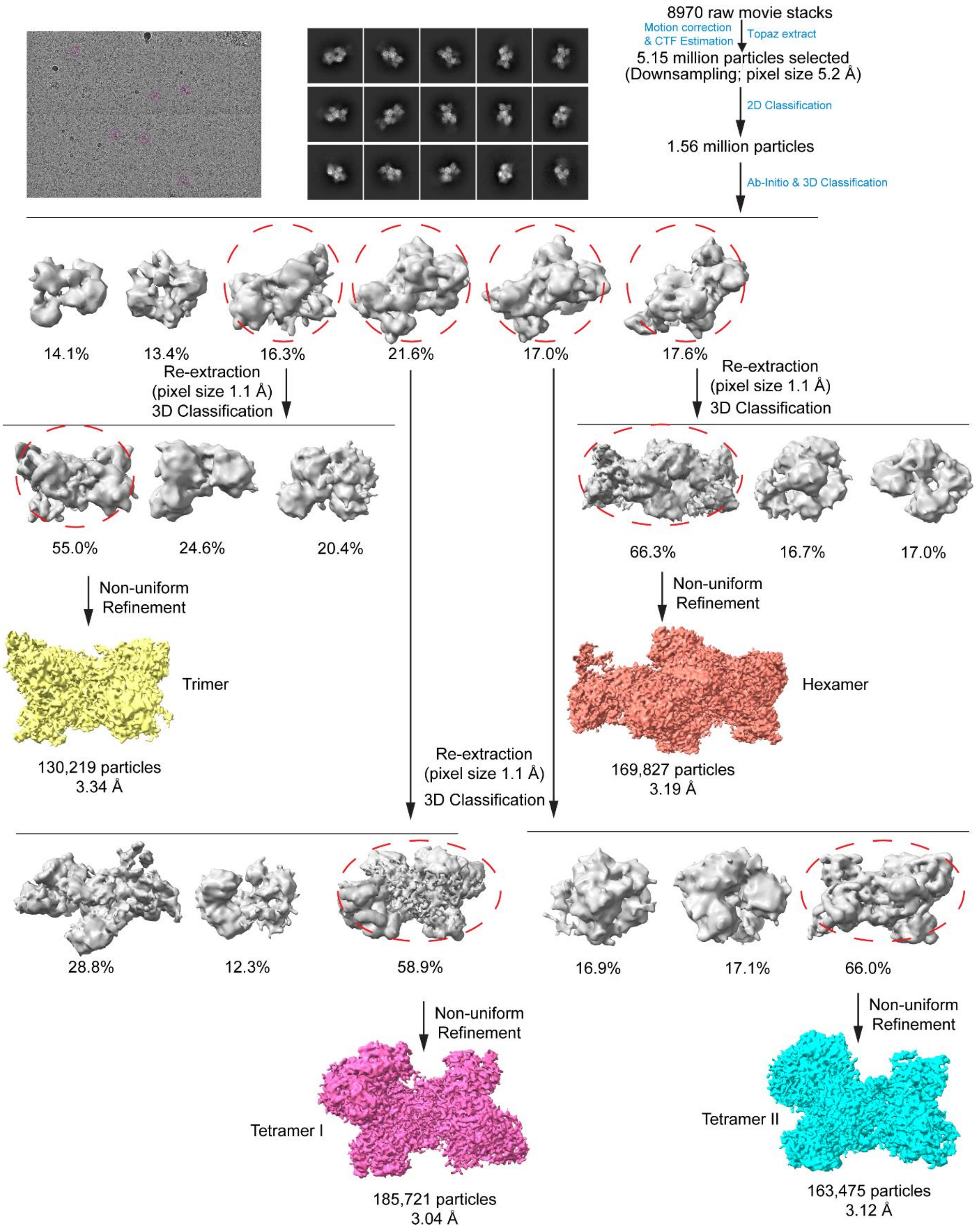


**Figure S2. Data processing workflow for cryo-EM reconstruction of the DNMT3B assemblies.** Note that the particle images were initially downsampled to a pixel size of 5.2 Å. After 2D and 3D classifications, the particles corresponding to the four different assemblies of DNMT3B were re-extracted with a pixel size of 1.1 Å. The final cryo-EM maps after cryoSPARC refinement were sharpened with a B-factor value of -92 $Å^2$, -84 $Å^2$, -91 $Å^2$ and -88 $Å^2$, respectively.

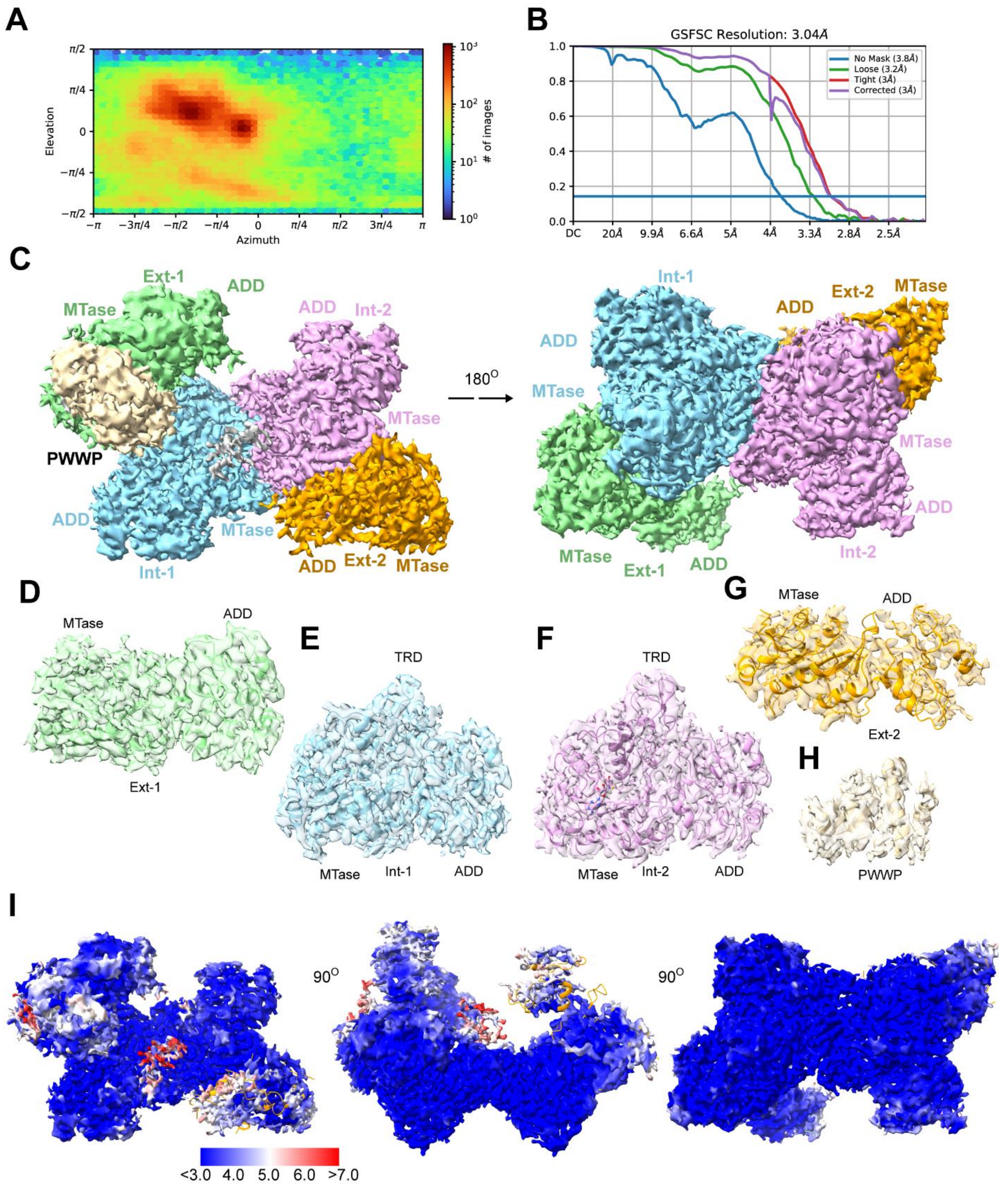


**Figure S3. Cryo-EM reconstruction of DNMT3B tetramer I.** (A) Orientation distribution map of DNMT3B tetramer I. (B) Fourier shell correlation (FSC) curve of DNMT3B tetramer I map as a function of resolution using cryoSPARC output. (C) Final density map of DNMT3B tetramer I in two opposite diections. The Ext-1, Int-1,

Int-2 and Ext-2 subunits, and the PWWP domain are colored in green, cyan, pink, orange and wheat, respectively. The unmodeled density corresponding to the N-terminal regions of DNMT3B subunits is colored in grey. (d-h) Strucutral model and associated density for the Ext-1 (D), Int-1 (E), Int-2 (F) and Ext-2 (G), with the ADD and MTase domains of each subunit labeled. (H) Strucutral model and associated density for the PWWP domain. (I) Local resolution map of DNMT3B tetramer I.

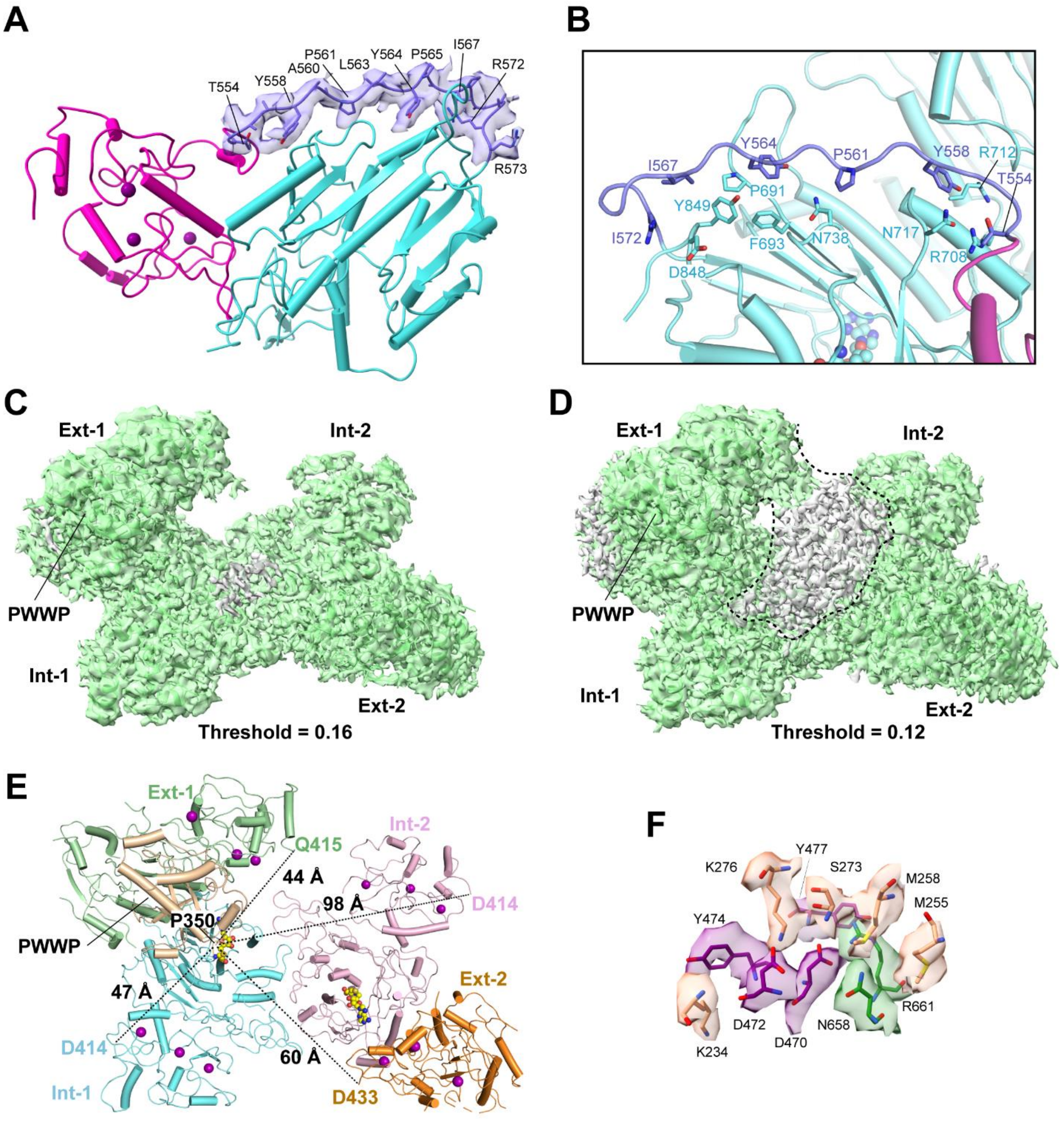


**Figure S4. Cryo-EM analysis of the N-terminal domains of DNMT3B tetramer I.** (A) Ribbon diagram of the Int-1 subunit, with the density associated with the ADD-MTase linker colored in slate. Select linker residues with traceable sidechain density are labeled. (B) Close-up view of the interaction between the ADD-MTase linker and MTase domain.

(C-C) Density map of the tetramer I at the threshold of 0.16 (C) and 0.12 (D). The unmodeled density is colored in grey and the rest in green. The unmodeled density and the N-terminus of the ADD domain of Ext-1 is surrounded by dashed line in (D). (E) The distances between the C-terminal residue P350 of the PWWP and the traceable N-terminal residue of the ADD domain of each subunit of the tetramer I. (F) Close-up view of the PWWP-ADD-MTase interface of the Ext-1 subunit, with the interacting residues of the PWWP, ADD and MTase domains colored in wheat, purple and green, respectively.

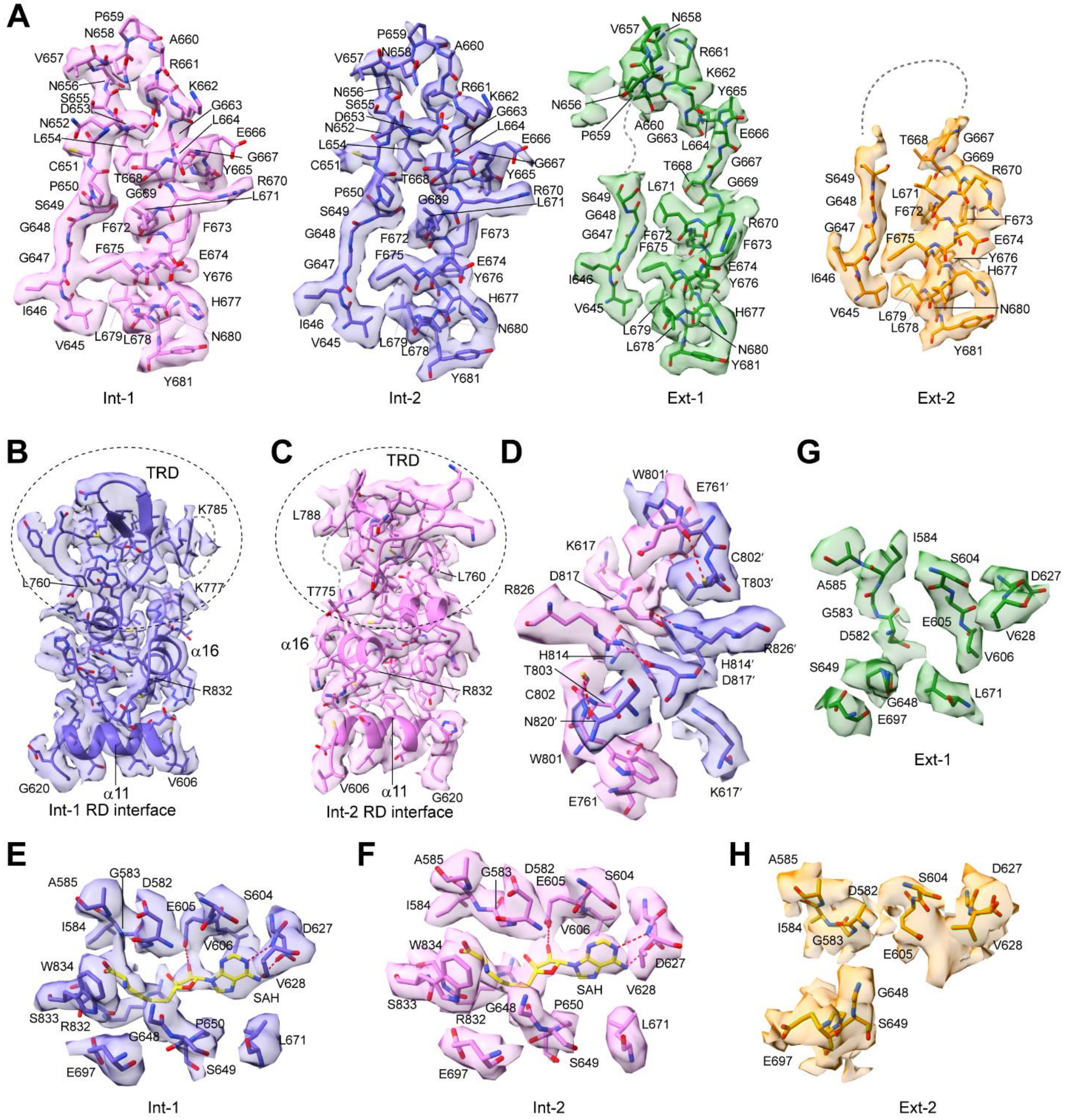


**Figure S5. Cryo-EM analysis of the oligomerization interfaces and cofactor-binding pocket of the tetramer I.** (A) The catalytic loop and its immediate downstream residues of the Int-1, Int-2, Ext-1 and Ext-2 subunits of the tetramer I. Note that the catalytic loop is well defined in Int-1 and Int-2 but becomes disordered to various extents in Ext-1 and Ext-2. (B-C) Structural elements (residues 760-832 and α11: 606-620) involved in the RD interface formation between the Int-1 (B) and Int-2 (C) subunits. The TRD of Int-1 or Int-2 is indicated by dashed circle. (D) Close-up view of the interacting residues at the RD interface between Int-1 (slate) and Int-2 (light magenta). The hydrogen bonds are shown as dashed lines. (E-H) The SAH-binding sites, formed in Int-1 (E) and Int-2 (F), are disrupted in Ext-1(G) and Ext-2 (H) of the tetramer I.

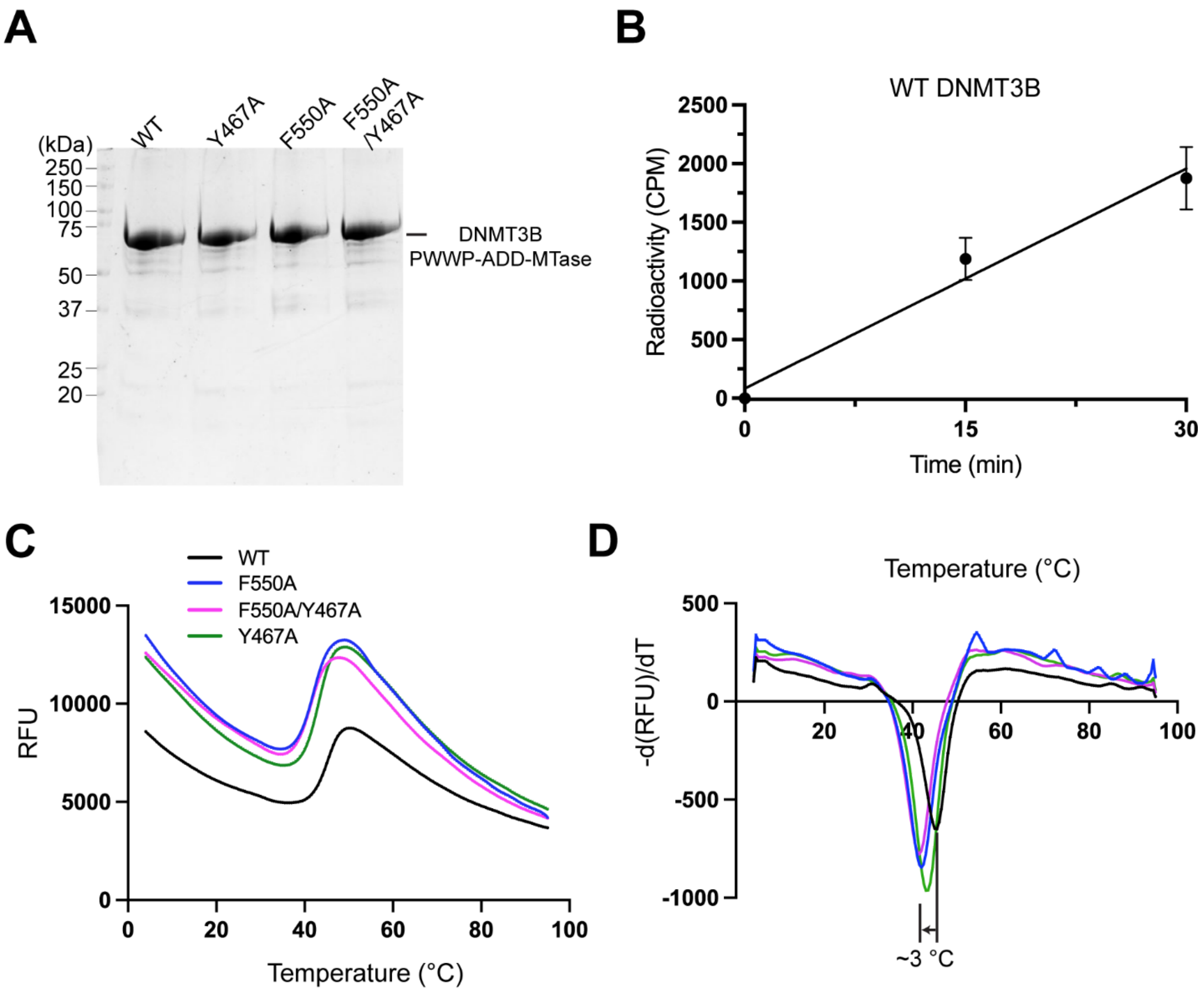


**Figure S6. Mutational analysis of the ADD-MTase interaction in DNMT3B.** (A) SDS-PAGE images of WT and mutant DNMT3B PWWP-ADD-MTase fragments. (B) *In vitro* DNA methylation kinetics of WT DNMT3B PWWP-ADD-MTase fragment. (C,D) Thermal shift assay for the WT and mutant DNMT3B PWWP-ADD-MTase fragment, with raw fluorescence data (C) and first derivative of the raw data (D) shown. The difference in melting temperature (ΔTm) between WT and F550A mutant is indicated.

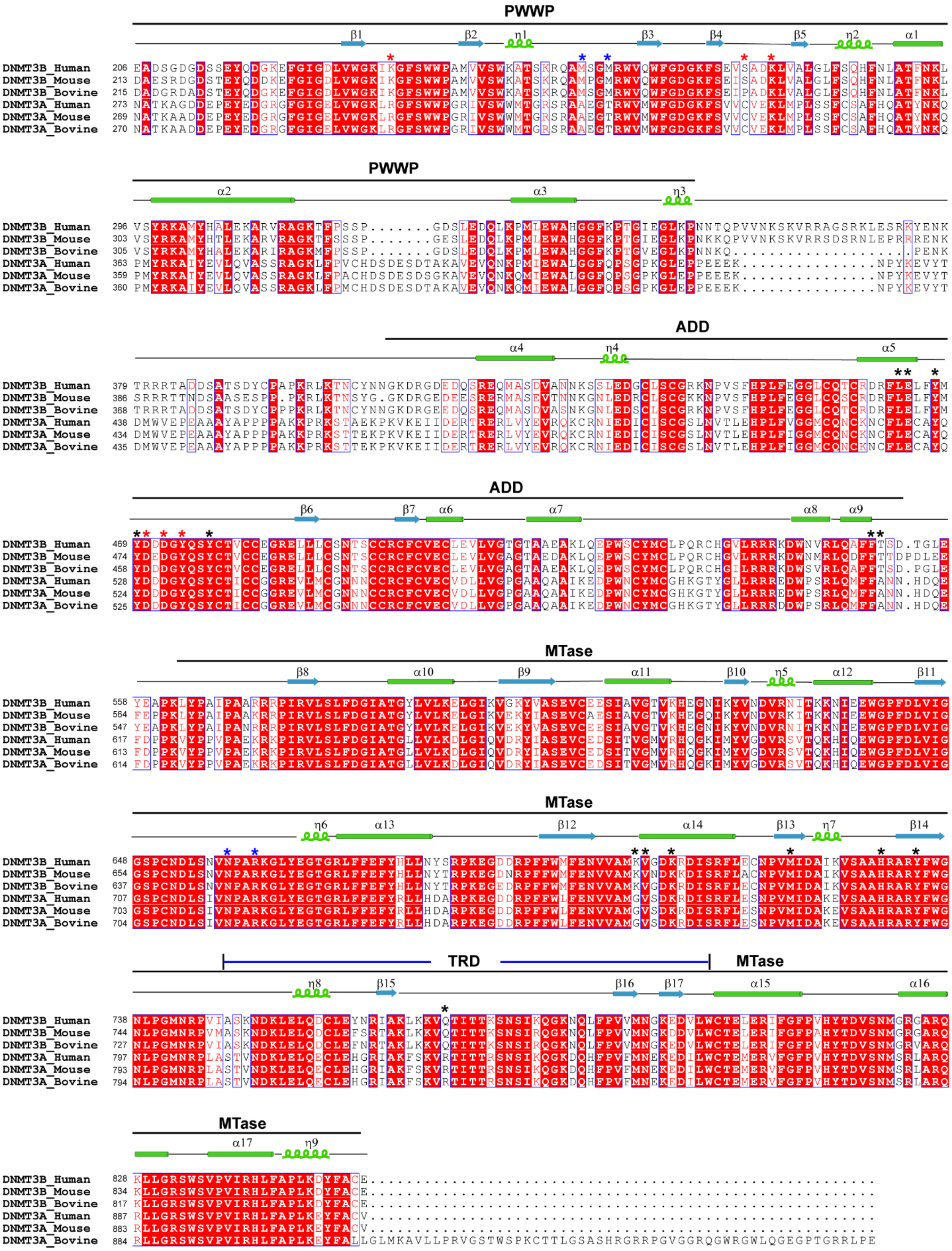


**Figure S7. Structure-based sequence alignment of DNMT3B and DNMT3A.** The aligned sequences encompass the PWWP, ADD and MTase domains of DNMT3B or DNMT3A from various species, with the secondary structures of DNMT3B indicated above. The TRD embedded in the MTase domain is marked. The DNMT3B residues involved in the PWWP-ADD, PWWP-MTase and ADD-MTase associations are marked with red, blue and black asterisks, respectively.

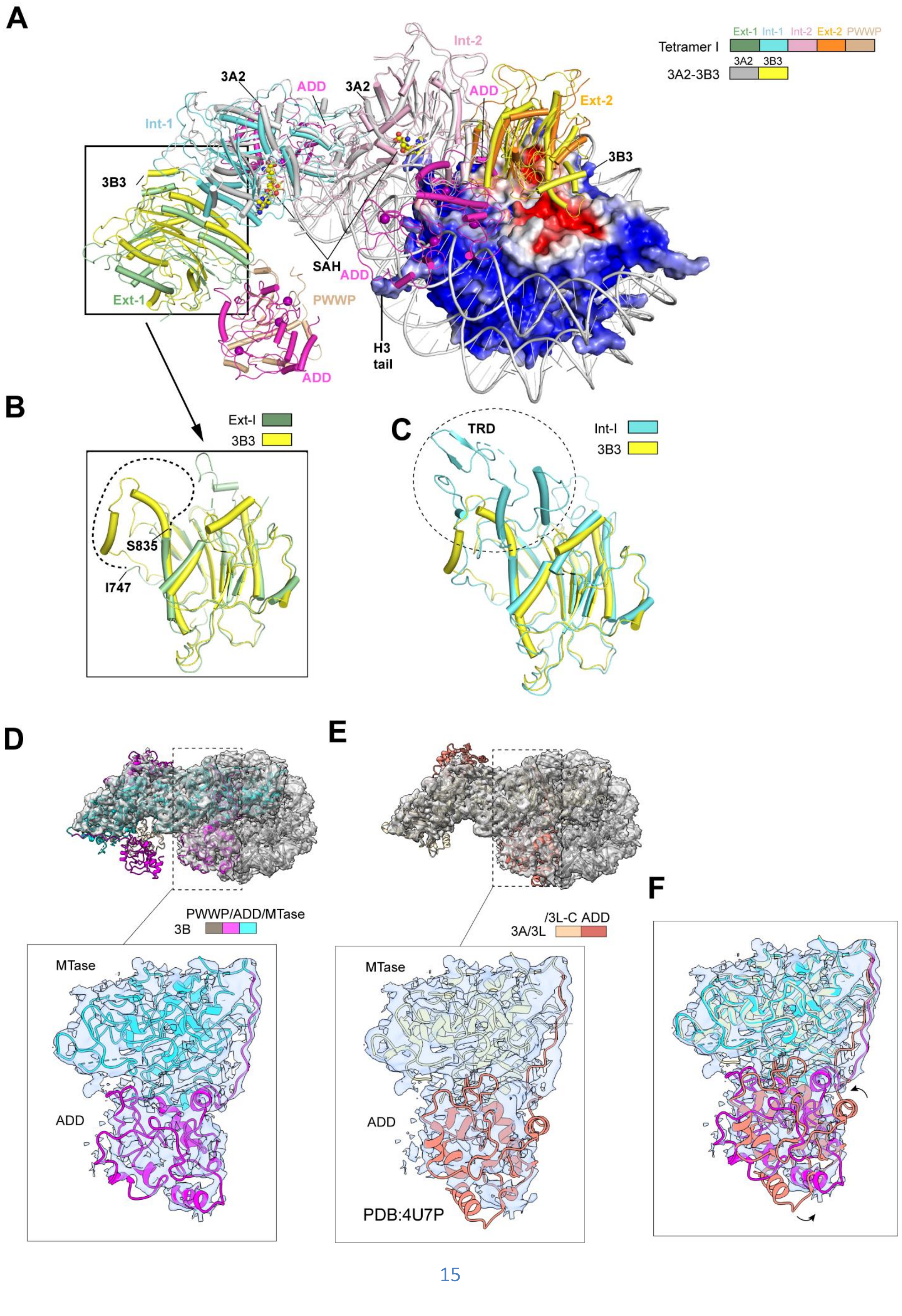
A
Tetramer I
Ext-1
Int-1
Int-2
Ext-2
PWWP
3A2-3B3
3A2
3B3
Int-2
3A2
ADD
3A2
ADD
Ext-2
Int-1
3B3
3B3
SAH
ADD
PWWP
Ext-1
H3
tail
ADD
B
Ext-I
3B3
S835
I747
C
TRD
Int-I
3B3
D
PWWP/ADD/MTase
3B
MTase
ADD
E
/3L-C ADD
3A/3L
MTase
ADD
PDB:4U7P
F

**Figure S8. Strucutral comaprison between DNMT3B tetramer I and the DNMT3A2-DNMT3B3-nucleosome complex.** (A) Structural overlay between DNMT3B tetramer I and the DNMT3A2-DNMT3B3 complex (PDB 6PA7). The DNMT3A, DNMT3B and DNMT3B3 fragments and nucleosomal DNA were shown in cartoon representation. Histone octamer is shown in electrostatic surface representation. (B) Structural alignment between the Ext-1 subunit of DNMT3B tetramer I and DNMT3B3 MTase domain, with the disordered segment of the Ext-1 shown by a dashed line. Note that residues 745-807 are missing in DNMT3B3 but present in DNMT3B1 (used in this study). For clarity, the ADD domain of the Ext-1 subunit is not shown. (C) Structural alignment between the Int-1 subunit of DNMT3B tetramer I and the MTase domain of DNMT3B3. The TRD, present in the Int-1 subunit but not in DNMT3B3, is indicated by dashed circle. (D,E) Docking the atomic model of DNMT3B tetramer I (D) and the DNMT3A-DNMT3L complex (PDB 4U7P) (E) each into the cryo-EM density map of DNMT3A2-DNMT3B3-NCP complex (EMD-20281). The relative positioning of the ADD-MTase domain of internal subunit II of the tetramer was shown in expanded views. Note that the DNMT3A ADD domain was defined but left unmodeled in the DNMT3A2-DNMT3B3-nucleosome complex (PDB 6PA7). Fitting of the DNMT3B ADD and DNMT3A ADD domains into the cryo-EM density yields model-map correlation coefficients of 0.48 and 0.29, respectively. (F) Overlap of the ADD-MTase domains of DNMT3B and DNMT3A in the density map of DNMT3A2-DNMT3B3-nucleosome complex. The repositioning of the ADD domain between DNMT3B and DNMT3A is indicated by the arrow.

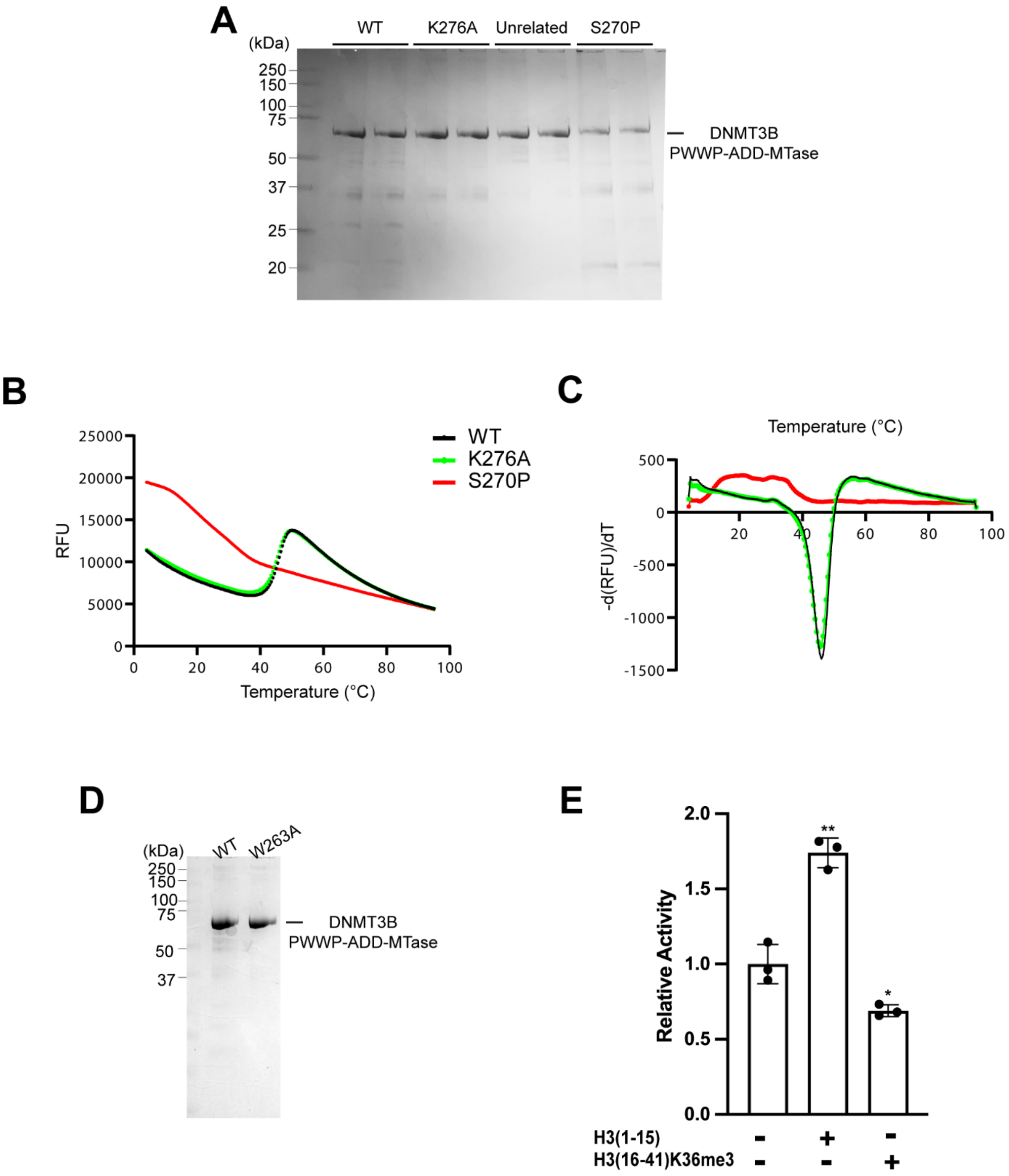
A
(kDa)
250
150
100
75
50
37
25
20
WT
K276A
Unrelated
S270P
DNMT3B
PWWP-ADD-MTase
B
RFU
25000
20000
15000
10000
5000
0
0
20
40
60
80
100
Temperature (°C)
WT
K276A
S270P
C
Temperature (°C)
-d(RFU)/dT
500
0
-500
-1000
-1500
20
40
60
80
100
D
(kDa)
WT
W263A
250
150
100
75
50
37
DNMT3B
PWWP-ADD-MTase
E
Relative Activity
2.0
1.5
1.0
0.5
0.0
**
*
H3(1-15)
H3(16-41)K36me3
- + -
- - +

**Figure S9. Biochemical analysis of the PWWP domain mutants of DNMT3B.** (A) SDS-PAGE analysis of the DNMT3B mutant. For enzymatic comparison, the relative amount of S270P and K276A over WT DNMT3B was adjusted based on their band intensities on SDS-PAGE. (B,C) Thermal shift assay for the WT and mutant DNMT3B PWWP-ADD-MTase fragments, with raw fluorescence data (B) and first derivative of the raw data (C) shown. (D) SDS-PAGE images of WT and W263A DNMT3B. (E) *In vitro* DNA methylation assay for W263A-mutated DNMT3B in the absence or presence of histone peptides. Data are mean ± s.d. (n = 3 biological replicates). The two-tailed Student t-tests were performed to compare peptide-free vs peptide-present. *, $p<0.05$; **, $p<0.01$.

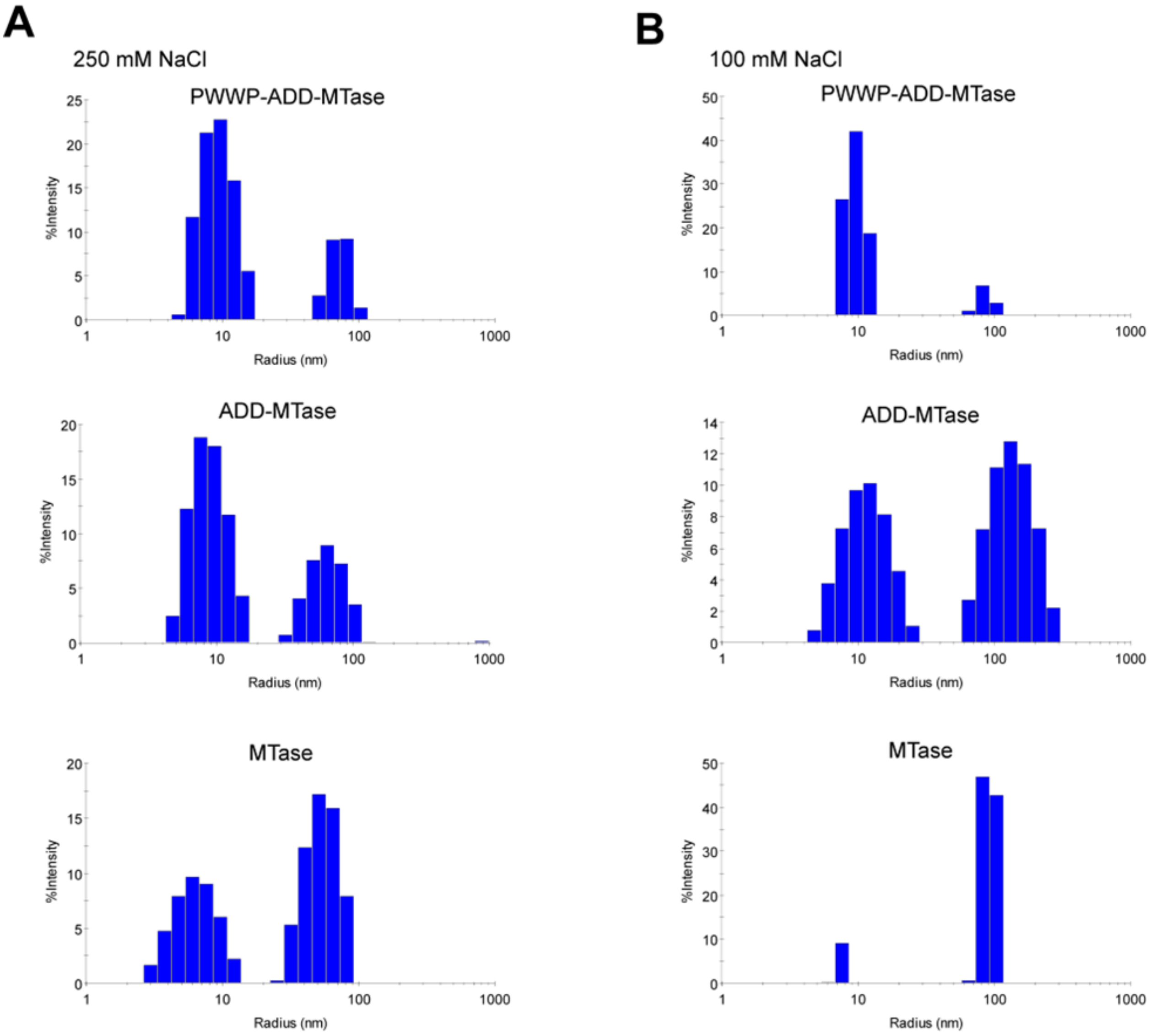


**Figure S10. DLS analysis of DNMT3B PWWP-ADD-MTase, ADD-MTase and MTase fragments.** (A, B) Comparison of the size distribution profile of DNMT3B PWWP-ADD-MTase, ADD-MTase and MTase fragments at 250 mM NaCl (A) and 100 mM NaCl (B). Under 250 mM NaCl salt condition, the size distribution profile of the PWWP-ADD-MTase fragment reveals a major low-molecular-weight population centered around hydrodynamic radius (Rh) of 9.6 nm and a minor high-molecular-weight population centered around Rh of 67.5 nm. In comparison, the ADD-MTase and MTase fragments exhibit more pronounced high-molecular-weight population. Reducing the salt concentration to 100 mM NaCl further strengthened the shift of the ADD-MTase and

MTase fragments, but not the PWWP-ADD-MTase fragment, toward the high-molecular-weight population.

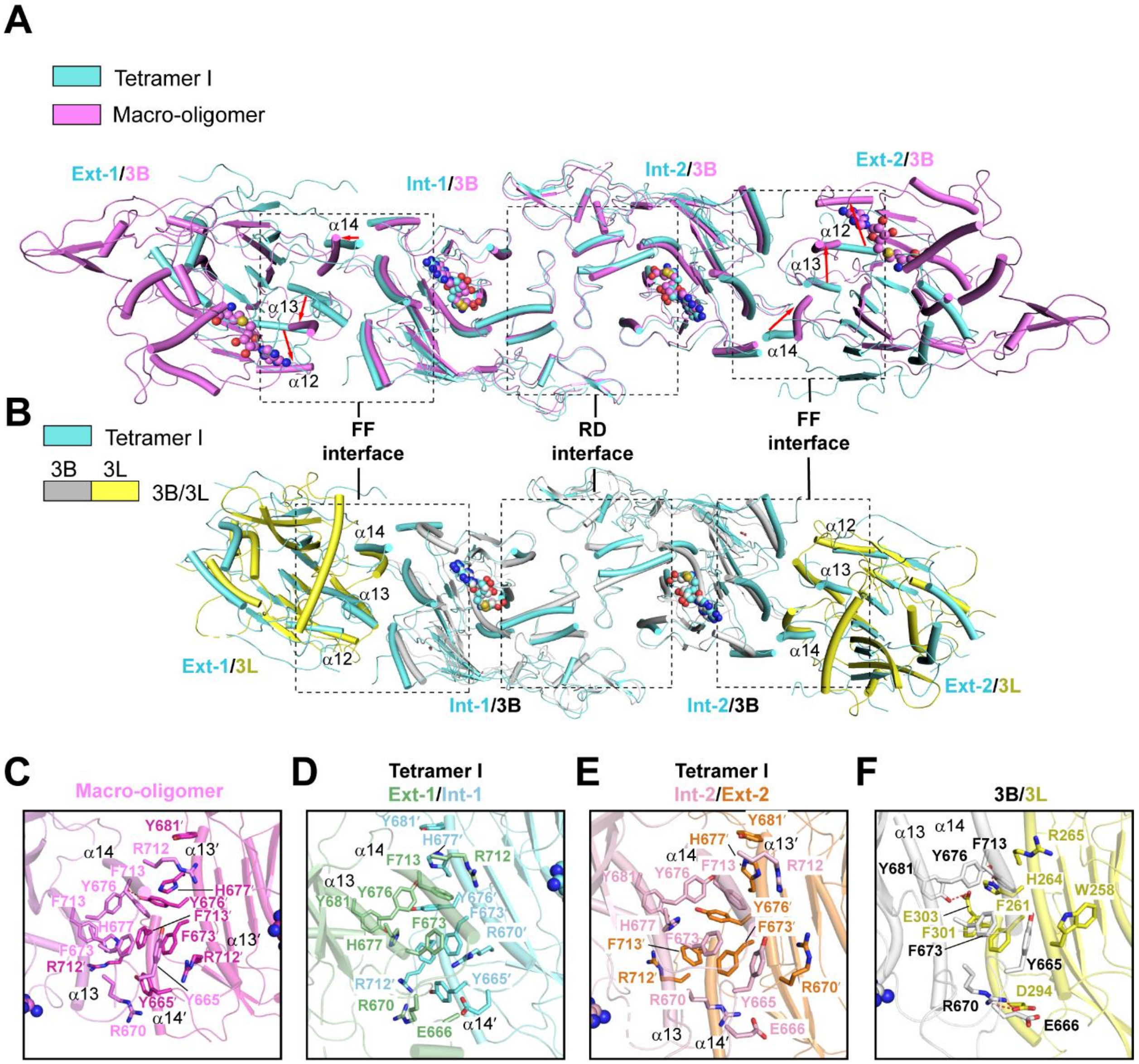


**Figure S11. Structural comparison of alternative DNMT3B assemblies.** (A,B) Structural alignment of DNMT3B tetramer I with the tetrameric unit of DNMT3B macro-oligomer (PDB 7V0E) (A) and the DNMT3B-DNMT3L heterotetramer (PDB 6KDP) (B), with the RD and FF interfaces indicated by dashed rectangles. The DNMT3B-DNMT3L complex shares a similar FF interface conformation as DNMT3B tetramer I, whereas the α-helices involved in the FF interface formation of the macro-oligomer show a lateral shift from those of DNMT3B tetramer I, indicated by red arrows. (C) Close-up view of the FF interface in the DNMT3B macro-oligomeric form. (D-E) Close-up view of the FF interface between the Ext-1 and Int-1 (D) and between the Int-2 and Ext-2 (E) subunits of the

DNMT3B tetramer I. (F) Close-up view of the FF interface between DNMT3B and DNMT3L in the DNMT3B-DNMT3L complex.

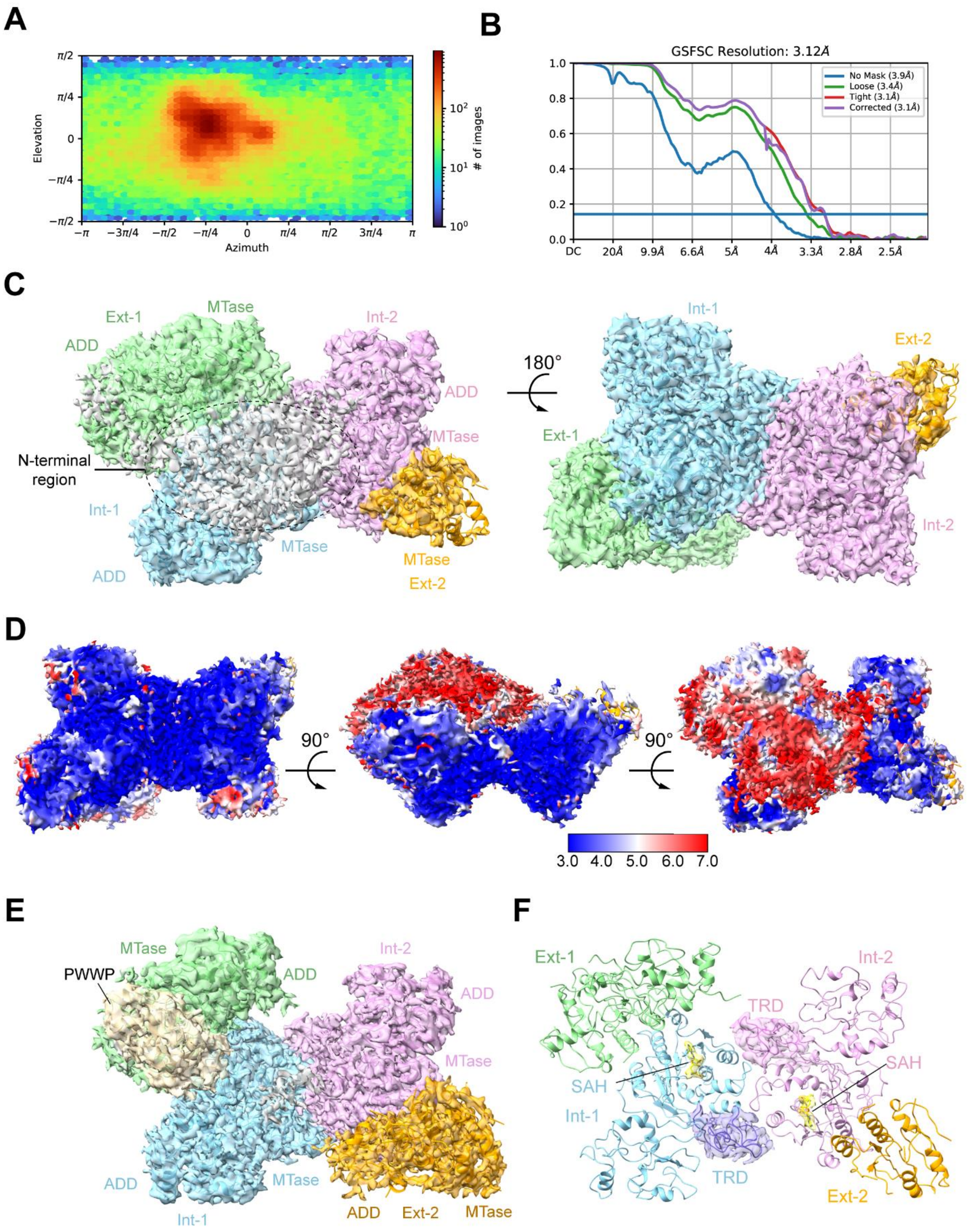


**Figure S12. Cryo-EM reconstruction of DNMT3B tetramer II.** (A) Orientation distribution map of DNMT3B tetramer II. (B) Fourier shell correlation (FSC) curve of

DNMT3B tetramer II map as a function of resolution using cryoSPARC output. (C) Final density map and strucutral model the of DNMT3B tetramer II in two opposite diections. The Ext-1, Int-1, Int-2 and Ext-2 subunits are colored in green, cyan, pink and orange, respectively. The unmodeled density corresponding to the N-terminal regions of DNMT3B is colored in grey. (D) Local resolution map of DNMT3B tetramer II. (E) Final density map and structural model of DNMT3B tetarmer I. The density threshold of (C) and (E) were adjusted to fit with the strucutral model. (F) Ribbon diagram of DNMT3B tetramer II, with the density for the TRD and SAH-binding sites of Int-1 and Int-2 shown.

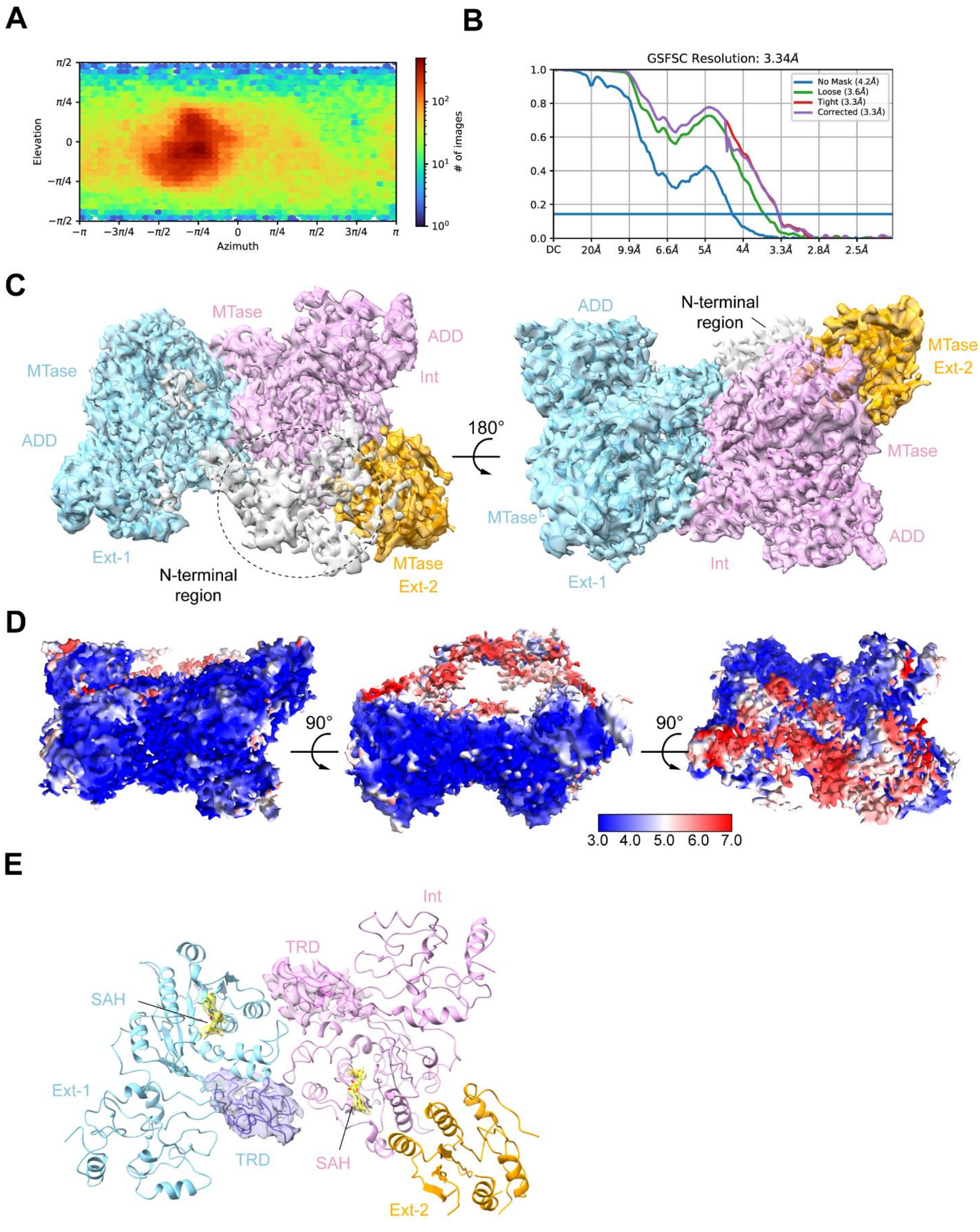
A
Elevation
Azimuth
### of images
B
GSFSC Resolution: 3.34Å
No Mask (4.2Å)
Loose (3.6Å)
Tight (3.3Å)
Corrected (3.3Å)
C
MTase
ADD
Int
MTase
ADD
Ext-1
N-terminal region
MTase Ext-2
180°
ADD
N-terminal region
MTase Ext-2
MTase
MTase
ADD
Int
Ext-1
D
90°
90°
3.0 4.0 5.0 6.0 7.0
E
Int
TRD
SAH
Ext-1
TRD
SAH
Ext-2

**Figure S13. Cryo-EM reconstruction of DNMT3B trimer.** (A) Orientation distribution map of DNMT3B trimer. (B) Fourier shell correlation (FSC) curve of DNMT3B trimer map as a function of resolution using cryoSPARC output. (C) Final density map and strucutral model the of DNMT3B trimer in two opposite diections. The Ext-1, Int and Ext-2 subunits are colored in cyan, pink and orange, respectively. The unmodeled density corresponding to the N-terminal regions of DNMT3B subunits is colored in grey. (D) Local resolution map of DNMT3B trimer. (E) Ribbon diagram of DNMT3B trimer, with the density for the TRD and SAH-binding sites of the Int and Ext-1 subunits shown.

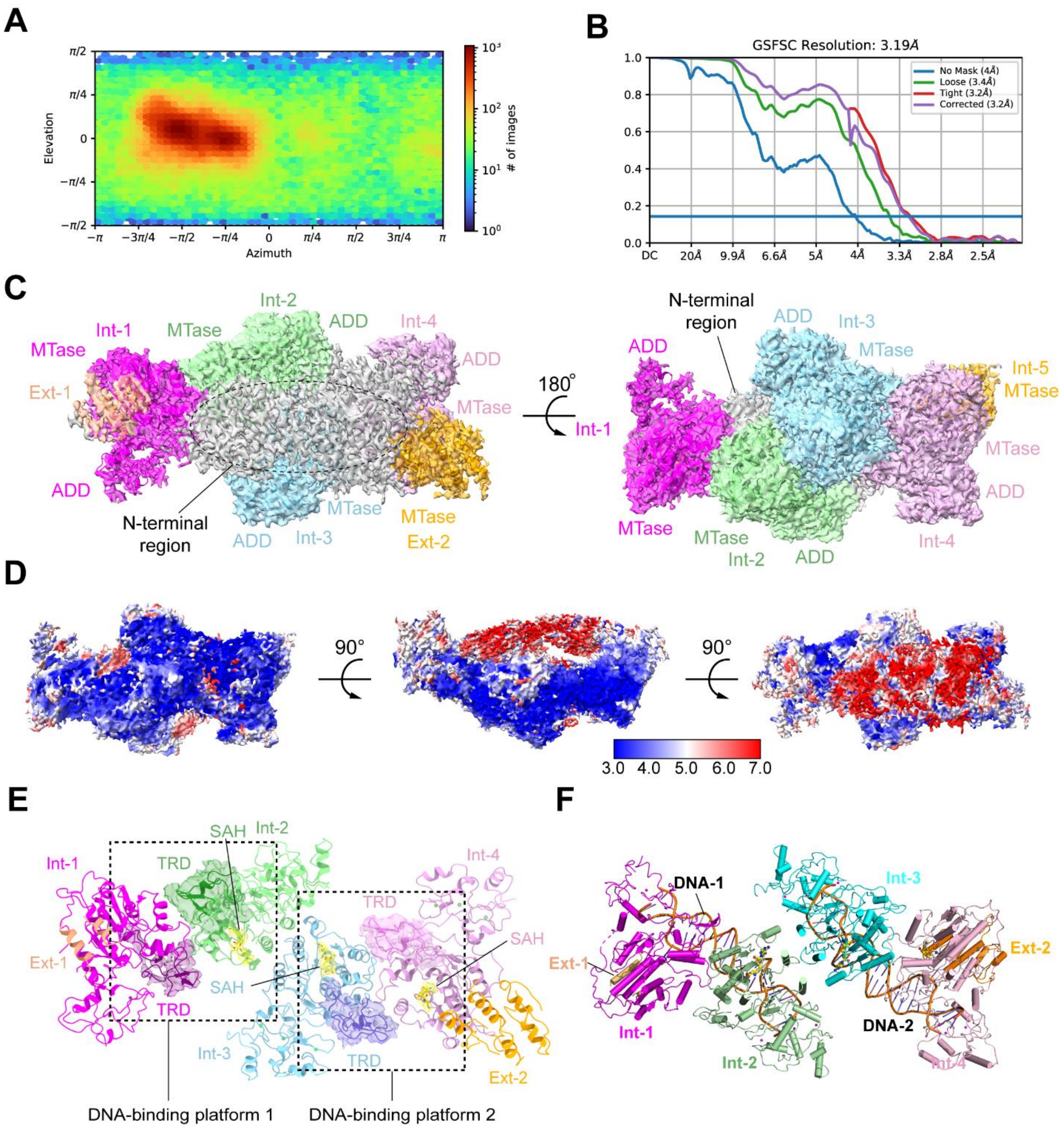


**Figure S14. Cryo-EM reconstruction of DNMT3B hexamer.** (A) Orientation distribution map of DNMT3B hexamer. (B) Fourier shell correlation (FSC) curve of DNMT3B hexamer map as a function of resolution using cryoSPARC output. (C) Final density map and strucutral model the of DNMT3B hexamer in two opposite diections. The Ext-1, Int-1, Int-2, Int-3, Int-4 and Ext-2 subunits are colored in salmon, magenta, green, cyan, pink and orange, respectively. The unmodeled density corresponding to the N-terminal regions of DNMT3B is colored in grey. (D) Local resolution map of DNMT3B trimer. (E) Ribbon diagram of DNMT3B hexamer, with the density for the TRD and/or SAH-binding sites of Int-1, Int-2, Int-3 and Int-4 shown. The hexamer forms two seperated DNA-binding platforms, which are marked by dashed rectangle. No SAH molecule is identified in the cofactor-binding pocket of Int-1, likely due to high conformational dynamcs of the corresponding region, as indicated by the density map in (C). (F) Structural model of the DNMT3B hexamer-DNA complex. The structural model was generated based on the DNA bound to DNMT3B-DNMT3L complex (PDB 6U8P).

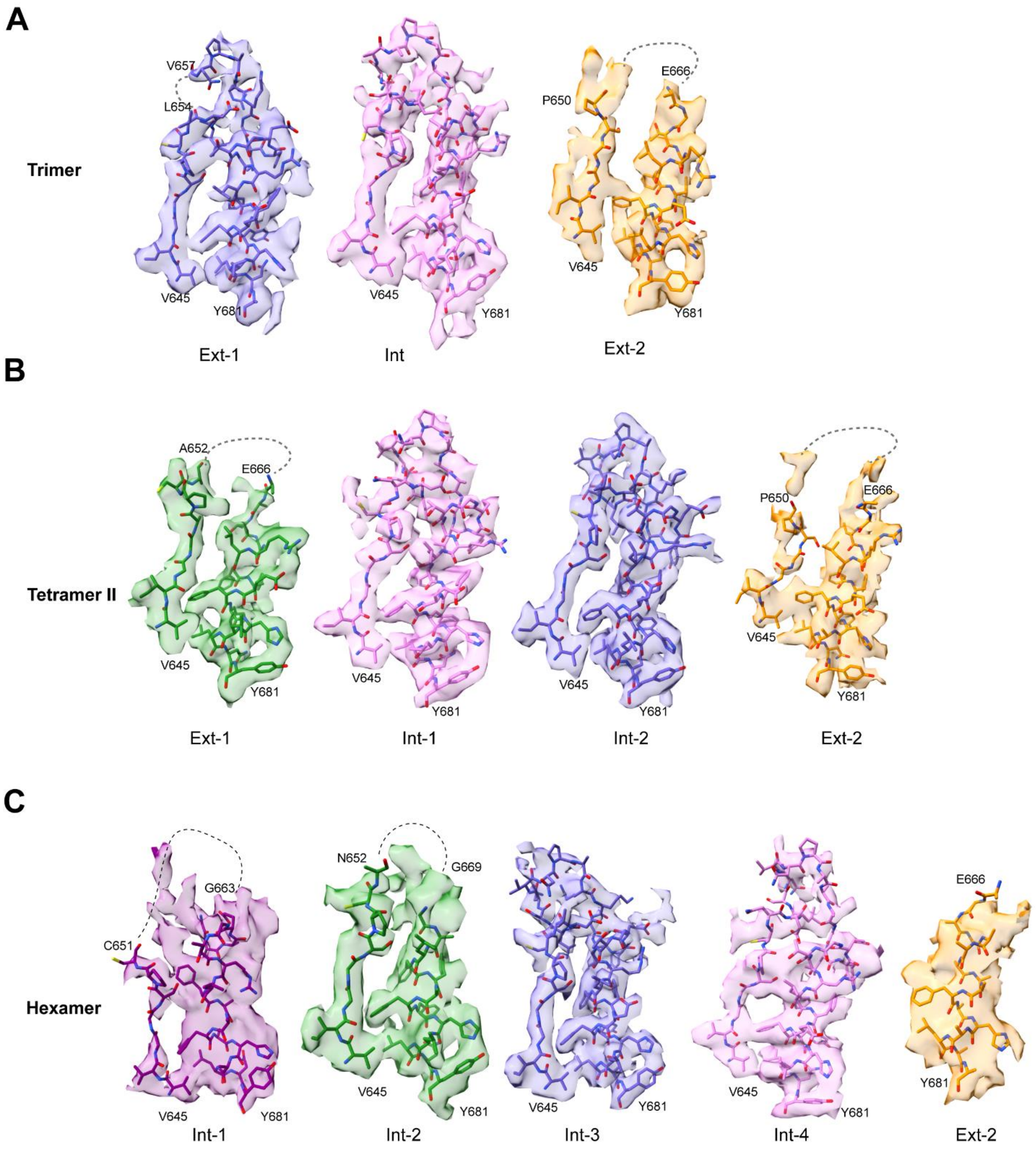


**Figure S15. Comparison of the catalytic loops among individual subunits in trimer, tetramer II and hexamer suggests a stablization effect by DNMT3B oligomerization.** (A) Catalytic loops and their immediate downstream residues of Ext-1, Int and Ext-2 subunits of DNMT3B tetramer. The disordered regions in those of Ext-1 and Ext-2 are shown as dashed lines. (B) Catalytic loops and their immediate downstream residues of Ext-1, Int and Ext-2 subunits of DNMT3B tetramer II. The disordered regions in those of Ext-1 and Ext-2 are shown as dashed lines. (C) Catalytic loops and their immediate downstream residues of Int-1, Int-2, Int-3, Int-4 and Ext-2 subunits of DNMT3B hexamer. The disordered regions in those of Int-1 and Int-2 are shown as dashed lines. Note that the catalytic loop is completely untraceable in Ext-1 and Ext-2.

#### Table S1. Cryo-EM data collection, structure refinement and validation statistics

| States<br><br>Codes | **Tetramer I** (EMD-28156, PDB 8EIH) | **Tetramer II** (EMD-28157, PDB 8EII) | **Trimer** (EMD-28158, PDB 8EIJ) | **Hexamer** (EMD-28159, PDB 8EIK) |
|---|---|---|---|---|
| **Data collection and processing** | | | | |
| Microscope | Titan Krios | | | |
| Camera | K3 BioQuantum | | | |
| Magnification | 81,000 | | | |
| Voltage (kV) | 300 | | | |
| Defocus range (μm) | -1.5 ~ -2.1 | | | |
| Exposure time (s) | 4 | | | |
| Dose rate($e^{-}$/Å$^2$/s) | 12.5 | | | |
| Number of frames | 40 | | | |
| Pixel size (Å) | 1.1 | | | |
| Micrographs (no.) | 8,970 | | | |
| Initial particles (no.) | 5,145,319 | | | |
| Symmetry imposed | *C1* | *C1* | *C1* | *C1* |
| Initial particles (no.) | 315,317 | 247,961 | 236,891 | 256,332 |
| Final particles (no.) | 185,721 | 163,475 | 130,219 | 169,827 |
| Map resolution (Å) | 3.04 | 3.12 | 3.34 | 3.19 |
| FSC threshold | 0.143 | 0.143 | 0.143 | 0.143 |
| **Refinement** | | | | |
| Initial model used | 5CIU, 7V0E, 4U7P | 7V0E, 4U7P | 7V0E, 4U7P | 7V0E, 4U7P |
| Model resolution (Å) | 3.2 | 3.4 | 3.7 | 3.6 |
| FSC threshold | 0.5 | 0.5 | 0.5 | 0.5 |
| Map sharpening *B* factor (Å$^2$) | -92 | -84 | -91 | -88 |
| Model composition | | | | |
| Non-hydrogen atoms | 11,622 | 10,399 | 7,761 | 13,409 |
| Protein residues | 1,562 | 1,364 | 1015 | 1806 |
| Ion (zinc) | 12 | 9 | 3 | 12 |
| SAH | 2 | 2 | 2 | 3 |
| *B* factors (Å$^2$) | | | | |
| Protein | 79.41 | 71.32 | 103.37 | 102.19 |
| Ion (zinc) | 181.63 | 161.26 | 195.44 | 205.38 |
| SAH | 39.43 | 40.42 | 50.54 | 44.57 |
| R.m.s. deviations | | | | |
| Bond lengths (Å) | 0.003 | 0.003 | 0.003 | 0.003 |
| Bond angles (°) | 0.573 | 0.670 | 0.662 | 0.633 |
| **Validation** | | | | |
| MolProbity score | 1.55 | 1.89 | 1.88 | 1.89 |
| Clashscore | 5.46 | 9.34 | 9.44 | 8.72 |
| Poor rotamers (%) | 0.27 | 0.1 | 0.38 | 0.39 |
| Ramachandran plot | | | | |
| Favored (%) | 95.22 | 94.25 | 94.38 | 93.60 |
| Allowed (%) | 4.78 | 5.75 | 5.62 | 6.40 |
| Disallowed (%) | 0 | 0 | 0 | 0 |

#### Table S2. ITC binding parameters

| PWWP | ADD | $K_d$ (μM) | N value | ΔH (cal/mol) | ΔS (cal/mol/deg) |
|---|---|---|---|---|---|
| WT | WT | 49.0 ± 3.5 | 1.06 ± 0.03 | 468 ± 20 | 21.4 |
| K276A | WT | 139.2 ± 0.5 | 1.01 ± 0.15 | 554± 125 | 19.5 |
| S270P | WT | NDB | | | |
| WT | D470A/D472A | 93.8 ± 9.7 | 0.97 ± 0.07 | 372 ± 43 | 19.6 |

The mean and S.D. were derived from two-independent measurements. NDB, no detectable binding.

#### References


1. L. Gao *et al.*, Structure of DNMT3B homo-oligomer reveals vulnerability to impairment by ICF mutations. *Nature communications* **13**, 4249.
